# IMPDH1 retinal variants control filament architecture to tune allosteric regulation

**DOI:** 10.1101/2021.08.03.454821

**Authors:** Anika L Burrell, Chuankai Nie, Meerit Said, Jacqueline C Simonet, David Fernández-Justel, Matthew C Johnson, Joel Quispe, Rubén M Buey, Jeffrey R Peterson, Justin M Kollman

## Abstract

IMP dehydrogenase (IMPDH), a key regulatory enzyme in purine nucleotide biosynthesis, dynamically assembles filaments in response to changes in metabolic demand. Humans have two isoforms: IMPDH2 filaments reduce sensitivity to feedback inhibition by the downstream product GTP, while IMPDH1 assembly remains uncharacterized. IMPDH1 plays a unique role in retinal metabolism, and point mutants cause blindness and disrupt GTP regulation. Here, in a series of cryo-EM structures we show that IMPDH1 assembles polymorphic filaments with different assembly interfaces in active and inhibited states. Retina-specific splice variants introduce structural elements that reduce sensitivity to GTP inhibition, including stabilization of the active filament form. Finally, we show that IMPDH1 disease mutations fall into two classes: one disrupts GTP regulation and the other has no in vitro phenotype. These findings provide a foundation for understanding the role of IMPDH1 in retinal function and disease and demonstrate the diverse mechanisms by which metabolic enzyme filaments are allosterically regulated.

## INTRODUCTION

Cells have evolved a myriad of ways to maintain precise and balanced pools of purine nucleotides. Purines are essential components of RNA and DNA, provide energy and act as cofactors for many enzymatic reactions. Maintaining a balance between purine pools is necessary to cell survival. In most tissues, a complex and highly regulated interplay between salvage and de novo biosynthesis pathways maintains optimal nucleotide concentrations. In cells with high purine demands, like proliferating cells, both pathways are upregulated^1^.

IMP dehydrogenase (IMPDH) is a highly conserved enzyme that catalyzes the first committed step in GTP synthesis. IMPDH sits at a critical branch point between adenine and guanine nucleotide synthesis, where its regulation is critical for balancing flux through the two pathways (Fig. 1a). In vertebrate cells, IMPDH forms filamentous ultrastructures in response to high demand for guanine nucleotides^2–4^. Humans express two isoforms of IMPDH that have 84% sequence identity. Much of the research on human IMPDH has been focused on IMPDH2 because it plays a critical role in the immune response and is upregulated in proliferating cells^5–7^;for example, activation of T-cells drives assembly of IMPDH2 into filaments ^8,9^. In vitro studies have shown that incorporation of IMPDH2 into filaments prevents the enzyme from adopting a fully inhibited conformation, providing an additional layer of regulation that decreases GTP feedback inhibition^10^. IMPDH1 also assembles filaments^11^, but whether polymerization plays a similar role in regulation of this isoform has been unclear. Understanding the mechanisms of IMPDH1 regulation is particularly important, as point mutations in humans lead to retinal degeneration of varying severity, pointing to a key role of IMPDH1 in retinal metabolism^12^.

**Fig 1.**
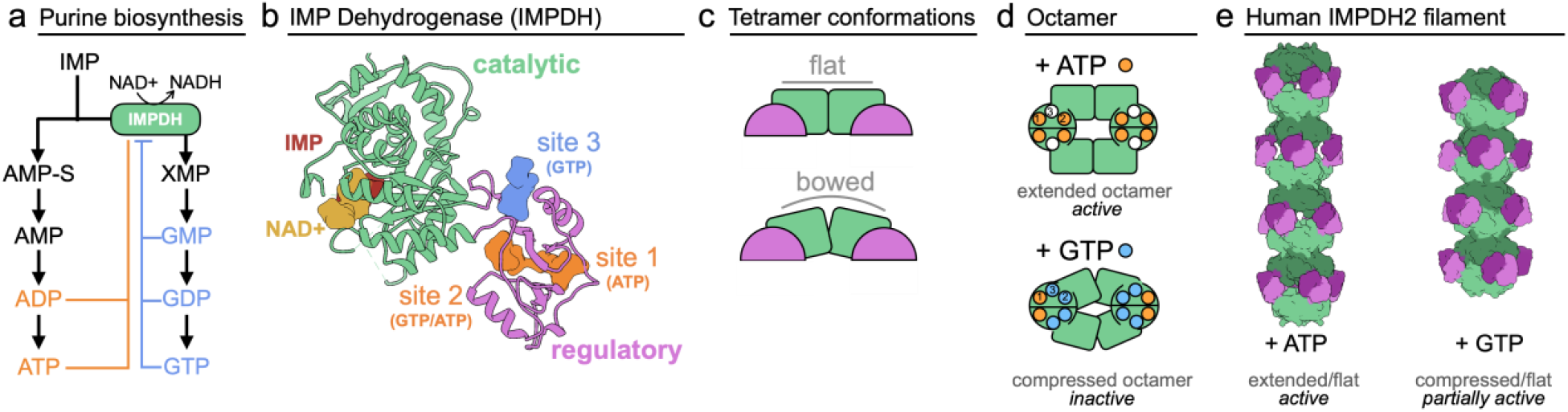
IMPDH structure and function. **a**, Purine biosynthesis pathway. **b**, IMPDH monomer (6u9o) has a catalytic domain (green) that binds IMP and NAD+ in the active site, and a regulatory domain (pink) with three allosteric nucleotide binding sites. **c**, IMPDH is a tetramer in solution and can adopt a flat or bowed conformation. Side view of tetramers are depicted, so that only two monomers are visible. **d**, ATP (sites 1&2) or GTP (sites 2&3) binding promotes octamer assembly. **e**, IMPDH2 octamers can assemble into filaments of stacked octamers.

IMPDH quaternary structure is directly linked to activity and regulation by adenine and guanine nucleotides. The monomer is composed of a catalytic and a regulatory cystathionine β-synthase domain, and constitutively assembles into tetramers through interactions of the catalytic domains^13^ (Fig. 1b,c). IMP is converted to xanthosine monophosphate (XMP) in an NAD+ dependent reaction in the catalytic domain that has been extensively characterized^14^. The regulatory domain has three allosteric nucleotide binding sites that bind adenine and guanine nucleotides^11,15,16^ (Fig. 1b). Site 1 has a preference for ATP/ADP, site 2 binds ATP/ADP and GTP/GDP competitively, and site 3 exclusively binds to GTP/GDP. Binding of nucleotides in sites 1 and 2 drives dimerization of the regulatory domains and formation of octamers, and the balance between ATP and GTP binding dictates whether octamers adopt an extended/active conformation or a GTP-bound compressed/inactive conformation (Fig. 1d)^11,17^. GTP binding at both competitive site 2 and the GTP-only site 3 induces two key conformational changes in IMPDH2: compression of the octamer and flexing of the catalytic domains from an active “flat” to an inactive “bowed” conformation (Fig. 1c,d). These changes inactivate IMPDH2 by preventing essential loop movements in the core of the enzyme^10,15,18^. Nucleotide-dependent assembly of IMPDH2 into filaments stabilizes the flat conformation (Fig. 1c), but does not prevent compression, allowing filamentous IMPDH2 to remain partially active even at high GTP concentrations, consistent with the role of IMPDH2 in expanding guanine nucleotide pools during proliferation^10,19^ (Fig. 1e).

Mutations in IMPDH1 lead to retinal degeneration in humans, a condition known as retinitis pigmentosa, but the molecular mechanism of disease remains unknown^12^. The activity of the mutant enzymes appears to be normal, but some mutations reduce sensitivity to GDP inhibition^11^. Despite expression of IMPDH1 in almost all tissues^20,21^ and the ubiquitous cellular need to maintain balanced purine pools, the retina appears to be the only affected tissue. This may be due to the very high and specific demands for ATP and cGMP in the retina ^22^. Photoreceptors have an unusually high demand for ATP, particularly in the dark^23^, and cyclic GMP is the key signaling molecule in the phototransduction cascade^24–26^. Furthermore, the retina is uniquely dependent on IMPDH1, because expression of both IMPDH2 and the key purine salvage enzyme HPRT are very low in the tissue^211^. The complex dependence of photoreceptor function on balanced purine production may lead to severe consequences if IMPDH1 is misregulated^27,28^.

In the retina, IMPDH1 is expressed as two splice variants that have additions to each terminus. In the IMPDH1(546) variant, five residues at the C-terminus of the canonical enzyme are replaced by 37 new residues, most of which are predicted to be unstructured. The other retinal variant IMPDH1(595) contains the same C-terminal extension, plus 49 residues at the N-terminus. Most of the N-terminal addition is predicted to be unstructured, except a short, predicted helix near the canonical N-terminus^21,29,30^ (Supp. Fig. 1). Mouse retinal splice variants have previously been reported to have reduced GTP inhibition compared to the canonical variant, but there has yet to be a structural explanation for the change^31^.

Here, we show that canonical human IMPDH1 assembles into filaments with different structures depending on its activity state. Active IMPDH1 filaments closely resemble active IMPDH2 filaments. The GTP-inhibited IMPDH1 filament, however, uses completely different assembly contacts, which allows the filament to accommodate a fully inhibited bent conformation of the enzyme. Thus, unlike IMPDH2, canonical IMPDH1 does not experience reduced feedback inhibition in the filament. The retinal variants, however, do have reduced sensitivity to GTP feedback, with independent mechanisms for the N- and C-terminal extensions in influencing IMPDH1 conformation. The N-terminal extension introduces specific structural changes in filament architecture that stabilize the partially active flat, compressed enzyme conformation, while the C-terminal extension likely reduces the stability of the compressed conformation. Finally, we characterize IMPDH1 retinopathy mutations in the retinal variants, revealing two distinct functional classes in vitro. Class I retinopathy mutants are not inhibited by GTP and do not adopt the compressed conformation, while Class II mutations appear identical to wildtype in biochemistry and filament assembly behavior.

## RESULTS

### IMPDH1 assembles into a filament in response to ATP or GTP binding

We tested the effects of the allosteric regulators ATP and GTP on IMPDH1 filament assembly and found broad similarities between IMPDH1 and IMPDH2. Using negative stain electron microscopy, we found that, like IMPDH2, IMPDH1 assembles into filaments of stacked octamers in the presence of ATP or GTP (Fig. 2a). The ATP-bound IMPDH1 filaments have a 110 Å rise and the GTP-bound IMPDH1 filaments have a 95 Å rise, consistent with extended/active and compressed/inactive octamers seen in IMPDH2 filaments^19^. We previously engineered a separation of function point mutation in IMPDH2 at the filament assembly interface, Tyr12Ala, which disrupts polymerization but does not affect enzyme activity^10,19^. The Y12A mutation also inhibits polymerization of IMPDH1, suggesting both isoforms assemble with similar interfaces (Fig. 2a).

**Fig 2.**
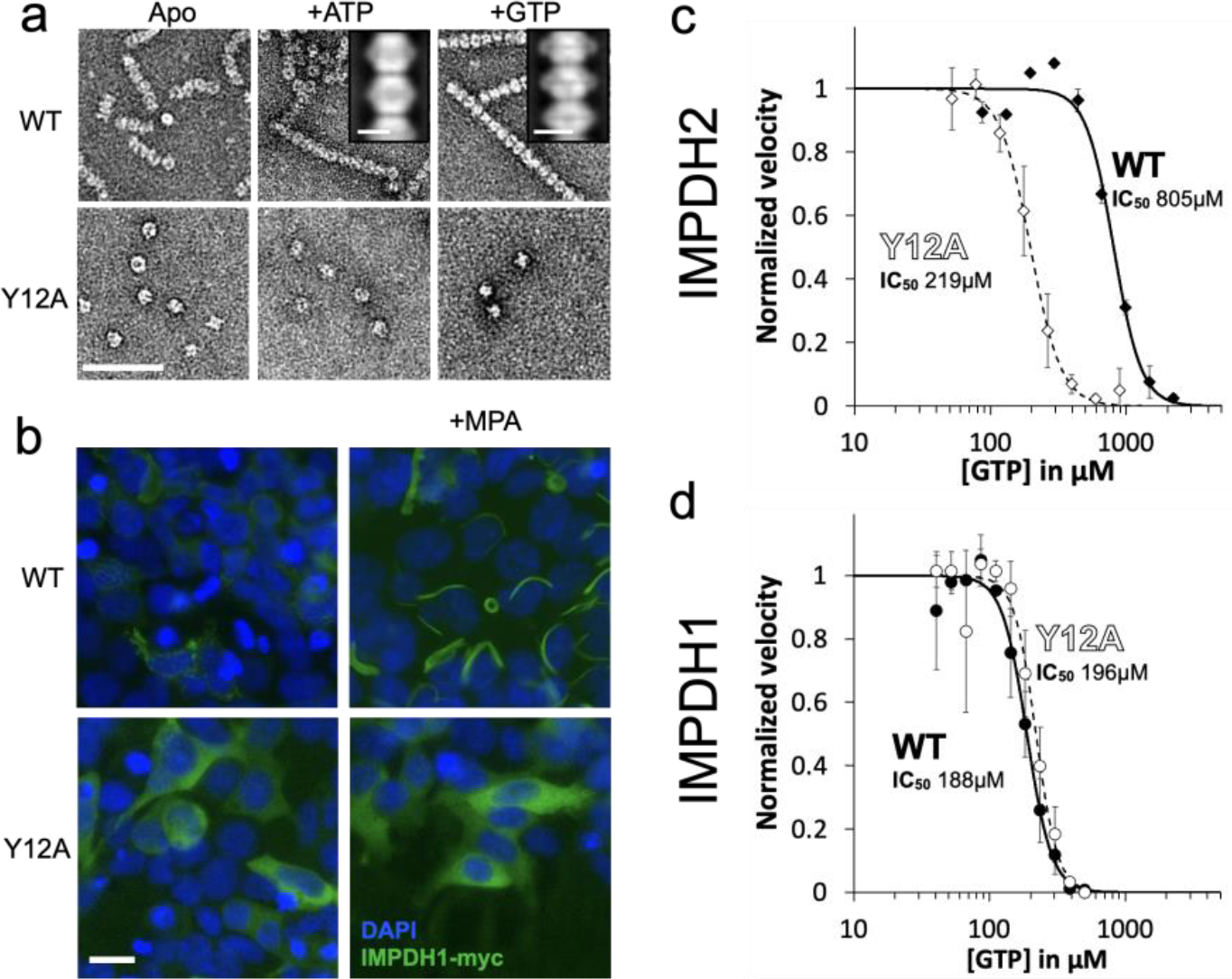
IMPDH1 assembles filaments and is sensitive to GTP inhibition. **a**, Negative stain EM of purified human IMPDH1. Addition of ATP or GTP promotes filament assembly of wildtype. Non-assembly mutation Y12A breaks both ATP- and GTP-dependent assembly. Insets are negative stain 2D class averages. Scale bar 100 nm. **b**, Anti-myc immunofluorescence of HEK293 cells transfected with IMPDH1-myc constructs (green). DAPI staining in blue. Cells were either left untreated or treated with 10 μM mycophenolic acid (MPA) to induce IMPDH2 filament assembly. Scale bar 20 μm. **c-d**, GTP inhibition curves of IMPDH2 or IMPDH1 WT (solid line) and the respective non-assembly Y12A protein (dashed line). Reactions performed with 1 μM protein, 1 mM ATP, 1 mM IMP, 300 μM NAD+, and varying GTP.

Previous studies have shown that IMPDH1 forms ultrastructures in cells, but the question remains whether the cellular ultrastructures assemble by the same mechanism as filaments in vitro^32^. To test this, we transiently transfected HEK293 cells with IMPDH1-myc, and induced filament assembly by treatment with the IMPDH inhibitor mycophenolic acid (MPA), a standard assay for cellular filament assembly^2,33,34^. In cells, IMPDH filaments appear to bundle together to assemble large ultrastructures several microns in length^35^. Staining with an anti-myc antibody shows strong induction of ultrastructure assembly with wildtype IMPDH1, but not with IMPDH1-Y12A, suggesting that the ultrastructures observed in cells are composed of filaments with an architecture similar to those we observe in vitro (Fig. 2b).

### IMPDH1 is more sensitive to GTP feedback inhibition than IMPDH2

We characterized the substrate kinetics of IMPDH1 and IMPDH2 and found them to be nearly identical (Supp. Table 1). However, we found a striking difference in sensitivity to GTP inhibition, with the GTP IC_50_ 4-fold lower for IMPDH1 than for IMPDH2 (Fig. 2c,d)^36^. Consistent with our prior results^10^, the non-assembly mutant IMPDH2-Y12A has a lower IC_50_ than the wildtype enzyme, but the same mutation has no effect on GTP affinity in IMPDH1 (Fig. 2c,d).

It is notable that IMPDH1-WT and Y12A mutants in both isoforms have similar GTP sensitivity and are fully inhibited at high GTP concentrations, while IMPDH2-WT retains basal activity even at GTP concentrations 6 times higher than the IC_50_ (Supp. Fig 2). This suggests conservation of the intrinsic allosteric regulation, with the difference being that polymerization reduces the inhibitory effect of GTP in IMPDH2 but appears to have no role in tuning the response of IMPDH1. This was surprising given the similarity of GTP-bound IMPDH1 and IMPDH2 filaments in our low-resolution negative stain imaging (Fig. 2a), so we next turned to higher resolution cryo-EM of IMPDH1 filaments to provide insight into differences in inhibition behavior.

### IMPDH1 assembles polymorphic filaments

We determined the structure of the IMPDH1 active filament bound to ATP, IMP, and NAD+ by cryo-EM, using the approach we developed to solve structures of IMPDH2 filaments, which have a rigid assembly interface between octamers but are more flexible within octamers^10^. This single-particle approach combines density subtraction, focused classification, and focused refinement of two different regions of IMPDH1 filaments: full octamers, and the assembly interface between stacked octamers (Supp. Fig. 3). This approach yielded octamer- and interface-focused reconstructions at 3.1 Å and 2.6 Å resolution, respectively (Fig. 3a–c) for active IMPDH1 filaments.

**Fig 3.**
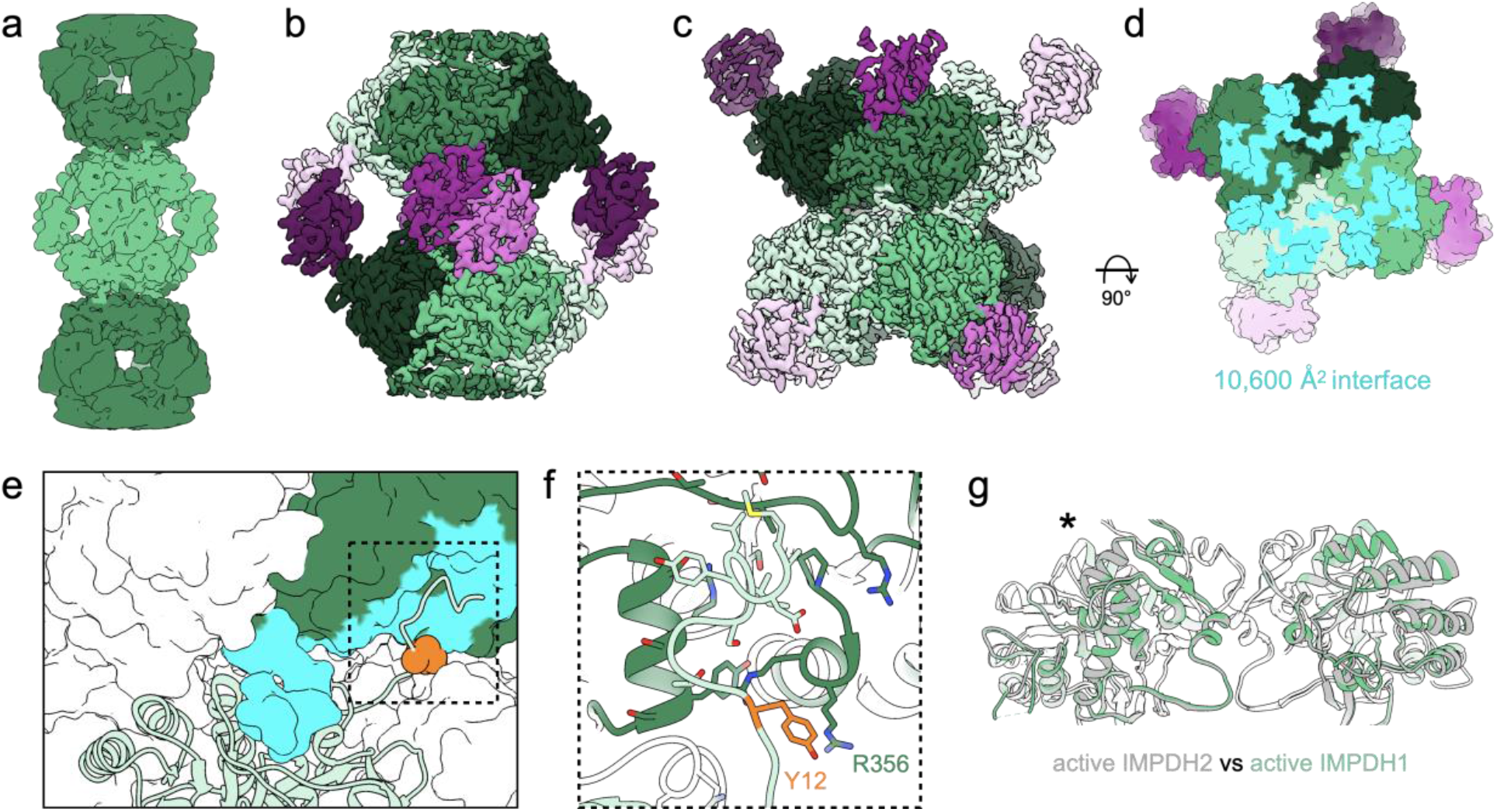
Structure of active IMPDH1 filaments (ATP/IMP/NAD+ bound). **a**, Low-pass filtered cryo-EM reconstruction colored by octamer. **b**, Octamer-centered single-particle reconstruction at Å, with catalytic domains in different shades of green and regulatory domains in shades of pink. **c**, Interface-centered single-particle reconstruction at 2.6 Å. **d**, Surface representation of the atomic model at the assembly interface, with buried residues in cyan. **e**, Surface representation of the filament interface with one monomer in ribbon (light green). Tyr12 is shown in orange spheres. The monomer it contacts across the octamer interface is green with residues forming the interface in cyan. **f**, Close-up ribbon view of the interface where Tyr12 in orange contacts R356 in the opposing monomer. **g**, Comparison of the catalytic tetramers of active IMPDH2 ATP/IMP/NAD+ (6u8s) (gray) to active IMPDH1 ATP/IMP/NAD+ (green). Aligned on monomers with asterisk.

The active state filament structure is highly conserved between IMPDH1 and IMPDH2. Like IMPDH2, the IMPDH1 active filament is composed of D4 symmetric stacked octamers with filament assembly contacts made between catalytic domains of opposing octamers. Each octamer has a rise of 113 Å and right-handed helical rotation of 30° between octamers. The interface buries a total of 10,600 Å^2^ (1,320 Å^2^ per monomer) (Fig. 3d) and is formed by residues 2-12 from the N-terminus of one monomer that sit in a groove between two helices in the catalytic domain of the opposing monomer (Fig. 3e,f). Tyr12, which breaks filament assembly when mutated to alanine, packs against arginine 356 on the opposing monomer (Fig. 3f). This filament architecture results in a tetramer in the flat conformation (Fig. 3g) similar to IMPDH2 (0.964 Å RMSD for alpha carbons over the catalytic domains of the tetramer) (Supp. Table 2). Overall, the filament interface and protomer conformations of IMPDH1 filaments are nearly identical to the IMPDH2 filament^10^.

The only major difference between isoforms in the active state filament appears to be the degree of flexibility in the octamer subunit. IMPDH2 in the active state shows extreme heterogeneity with mixed partially extended and partially compressed conformations within the same filament, due to flexibility between the catalytic and regulatory domains^10^. The interface-centered reconstruction reached a higher resolution than the octamer-focused reconstruction, likely due to some limited flexibility between domains. However, after extensive classification in the octamer-centered reconstruction we found that almost all protomers in the IMPDH1 filament structure are in the fully extended conformation. This may reflect a higher degree of cooperativity in the conformational state of IMPDH1 relative to IMPDH2.

We next solved a cryo-EM structure of the IMPDH1 inhibited filament bound to GTP, ATP, & IMP and found it assembles with a completely different architecture, with different interface contacts that lead to different helical symmetry (Fig. 4a–d; Supp. Video 1). The filament is still made up of D4 symmetric octamers but the N-terminal residues of one monomer now contact a different groove in the catalytic domain of the monomer in the opposing octamer (Fig. 4e). This change in interaction leads to a dramatic shift in helical geometry where the rotation between stacked octamers increases from 30° to 74°. The major contacts are between Tyr12 and Glu15 on the N-terminus and Glu487 and Lys489 on the opposing monomer (Fig. 4e,f). The involvement of Tyr12 in this interface explains why the Tyr12Ala point mutation also prevents assembly of this filament (Fig. 2a). The surface area buried by the interface is 75% smaller −2,800 Å^2^ buried at the octamer-octamer interface (350 Å^2^ per monomer) for the inhibited IMPDH1 filament compared to 10,600 Å^2^ for the active IMPDH1 interface (Figs. 3d, 4d). We refer to this interface as the “small interface” and the previously characterized interface in all IMPDH2 filaments and active IMPDH1 as the “large interface” (Supp. Video 1).

**Fig 4.**
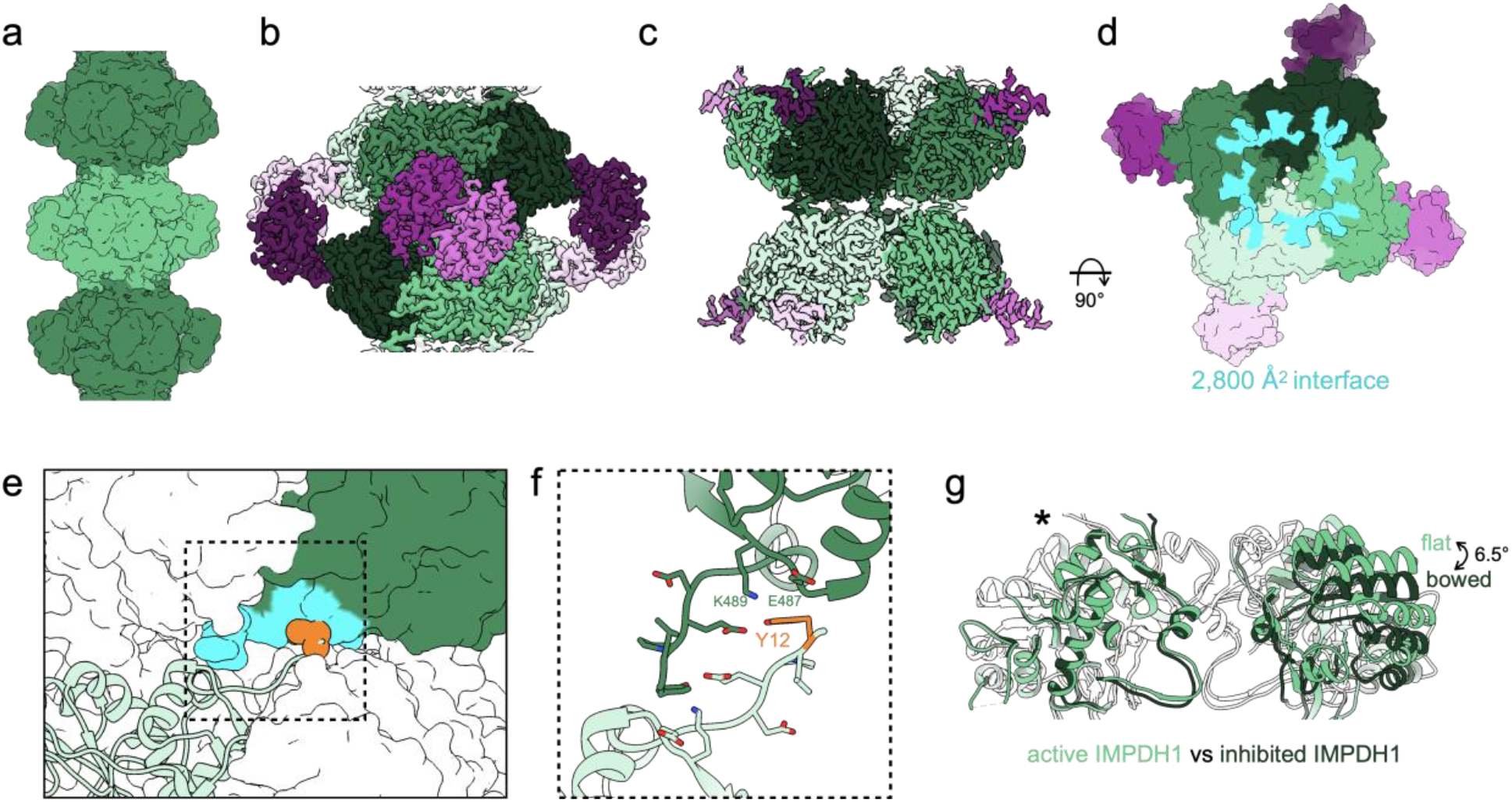
Inhibited IMPDH1 assembles with an alternate filament architecture (GTP/ATP/IMP bound) **a**, Low-pass filtered cryo-EM reconstruction colored by octamer. **b**, Octamer-centered single-particle reconstruction at 2.6 Å, with catalytic domains in different shades of green and regulatory domains in shades of pink. **c**, Interface-centered single-particle reconstruction at 2.6 Å. **d**, Surface representation of the atomic model at the assembly interface, with buried residues in cyan. **e**, Surface representation of filament interface with one monomer in ribbon (light green). Tyr12 is shown in orange spheres. The monomer it contacts across the octamer interface is green with residues forming the interface in aqua. **f**, Close-up ribbon view of the interface where Tyr12 in orange contacts Glu487 and Lys489 in the opposing monomer. **g**, Comparison of the catalytic tetramers of active IMPDH1 (light green) to inhibited IMPDH1 (dark green), show that the GTP-bound inhibited structure is in the bowed conformation. Aligned on monomers with asterisk.

The GTP-bound IMPDH1 filament accommodates a bowed tetramer that is fully inhibited (Fig. 4g). In contrast, GTP binding causes compression of IMPDH2, but filament contacts constrain the catalytic tetramers in a flat conformation, yielding a partially active flat, compressed state (Supp. Figs. 2a, 4a). A free IMPDH2 octamer is not restrained and able to adopt a completely inhibited bent, compressed state (Supp. Figs. 2a, 4a). GTP-boundIMPDH1 in filaments is nearly identical to the IMPDH2 free octamer bowed tetramer (Supp. Fig. 4 a,b) that is also fully inhibited (Supp. Fig. 2a). The conformational change can best be visualized by looking at protomers arranged diagonally to each other in the catalytic tetramer (chains A and C). Relative to the IMPDH1 active, flat conformation, there is a rotation of 6.5° between these two protomers in the inactive bowed conformation, resulting in an RMSD of 3.8 Å (Supp. Table 2). This finding provides an explanation for why IMPDH1 filaments don’t appear to directly affect regulation by GTP. For IMPDH1, GTP binding causes compression, but shifting filament assembly contacts also accommodate the bowed tetramer conformation, yielding a fully inhibited bowed, compressed state.

### IMPDH1 retinal variants assemble into filaments in response to ATP and GTP

We tested the effects of the allosteric regulators ATP and GTP on filament assembly of both IMPDH1 retinal variants (Fig. 5a) and found the response to be similar to canonical IMPDH1. Both retinal variants assemble filaments of stacked octamers in the presence of ATP or GTP, and IMPDH1(563) has a propensity to spontaneously assemble in the absence of ligands (Fig. 5b). ATP-bound IMPDH1 retinal variant filaments have ~110 Å rise and GTP-bound a ~95 Å rise, which is consistent with the IMPDH1 canonical filaments. In addition, we engineered a variant IMPDH1(563) that only has the N-terminal extension, to specifically test its effect on filament assembly, and it behaves like the retinal variants in negative stain EM. The mutation Tyr12Ala disrupts filament assembly in all the variants (Supp. Fig. 5).

**Fig 5.**
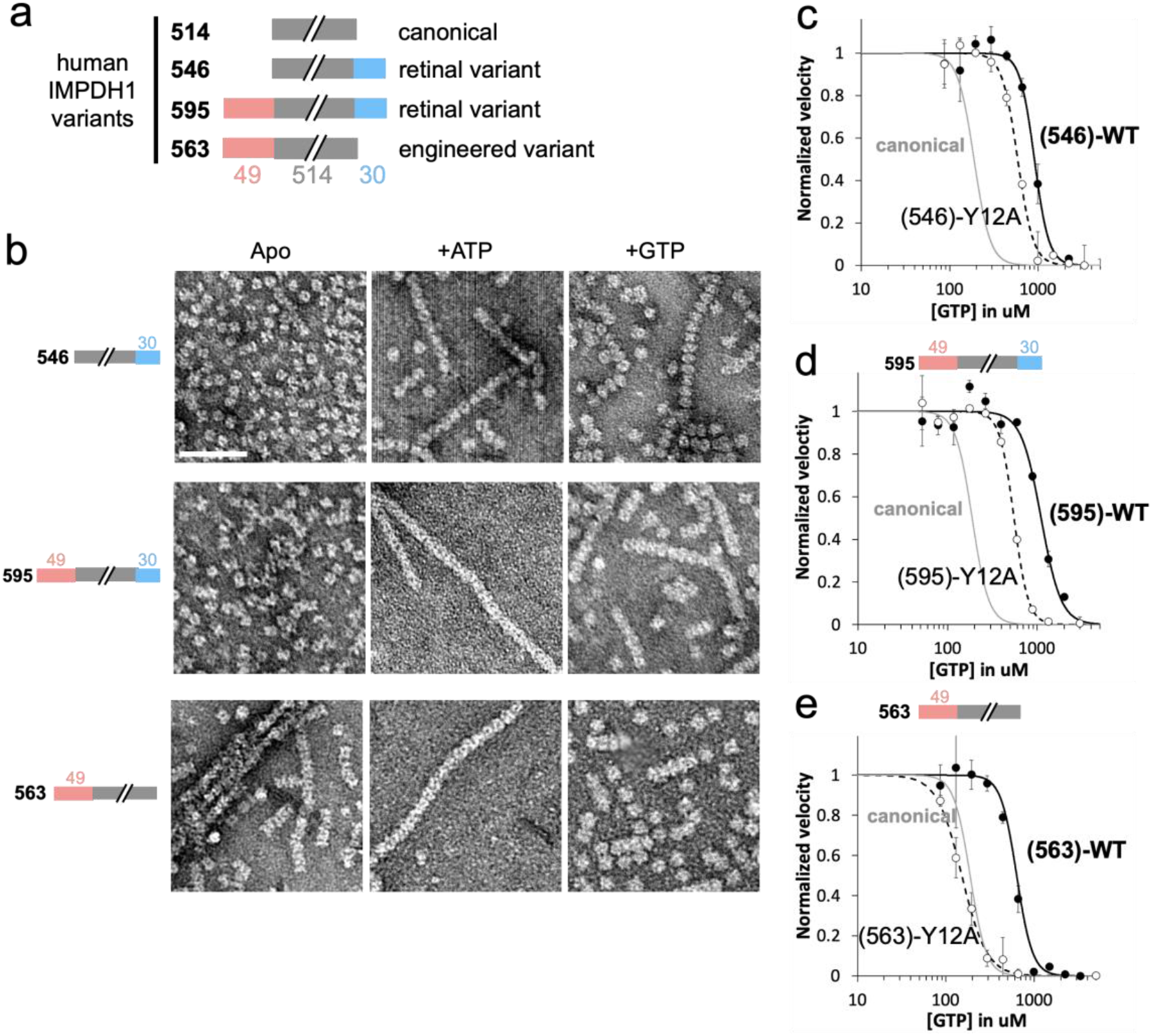
IMPDH1 retinal variants assemble filaments that resist GTP inhibition. **a**, Representation of IMPDH1 variant sequences. **b**, Negative stain EM of purified IMPDH1 variants in apo and nucleotide-bound states. Scale bar 100nm. **c-e** GTP inhibition curves of IMPDH1 variants (black solid line) and the respective non-assembly Y12A protein (black dashed line) compared to IMPDH1 canonical (solid gray line). **c**, Retinal variant IMPDH1(546). **d**, Retinal variant IMPDH1(595). **e**, Engineered variant IMPDH1(563). Reactions performed with 1 μM protein, 1 mM ATP, 1 mM IMP, 300 μM NAD+, and varying GTP.

### IMPDH1 retinal variants are less sensitive to GTP inhibition than canonical IMPDH1

The K_m_ values for IMP and NAD+ for the IMPDH1 retinal splice variants are very similar to each other and to the canonical variant (Supp Table 2). However, the retinal variants have higher IC_50_s for inhibition by GTP (Fig. 5c–e). IMPDH1(563) is 3 times less sensitive to GTP inhibition, IMPDH1(546) is almost 5 times less sensitive, and IMPDH1(595) 6 times less sensitive when compared to the canonical variant. These findings are consistent with previous results from the mouse IMPDH1 retinal splice variants^31^.

The sensitivity of IMPDH1 retinal variants and IMPDH2 to GTP is similar (Supp. Table 3); given the role of filament assembly in tuning GTP sensitivity in IMPDH2, we tested the effect of the Y12A non-assembly mutant on inhibition of IMPDH1 variants. All variants had increased sensitivity to GTP inhibition when the Y12A mutation was introduced (Fig. 5c–e). However, for IMPDH1(563), which only has the N-terminal extension, the GTP IC_50_ dropped to canonical levels, indicating that the effect of the N-terminal extension on IC_50_ is dependent on the ability to assemble filaments. Thus, the retina-specific N- and C-terminal additions appear to have distinct mechanisms that function independently to increase the GTP IC_50_.

### IMPDH1 retinal variants alter filament architecture

To gain insight into the mechanisms by which the splice variants alter GTP sensitivity, we determined structures of IMPDH1(595) and IMPDH1(546) in multiple ligand states by cryo-EM (Fig. 6a–c, Supp. Fig. 6). In the active state, both variants closely resemble active canonical filaments, with the enzyme in the fully extended, flat tetramer conformation, and the large assembly interface between octamers (Fig. 6a). In both active structures, most of the retinal C-term extension could not be resolved. Active canonical IMPDH1 can be resolved to residue 514, while active IMPDH1(595) could only be modeled to residue 504 and IMPDH1(546) to residue 515. In both IMPDH1(595) structures, there was clear density for a short alpha helix near the filament interface, composed of residues −22 to −4 (Fig. 6d). This helix packs between the catalytic core of the enzyme and the canonical N-terminal 15 residues that form the large interface (Fig. 6 e,f). In this position it also makes contacts with the neighboring protomer in the same tetramer with the major contact between residues Tyr-5 and His288 of the neighbor (Fig. 6g,h). The N-terminal end of the variant helix also contacts its symmetry mate on the monomer across the interface with Gln-21/Gln-15 (Fig. 6f). The helix appears to be positioned to stabilize the residues involved in the large interface filament interactions, and to stabilize the flat conformation of the catalytic tetramer.

**Fig 6.**
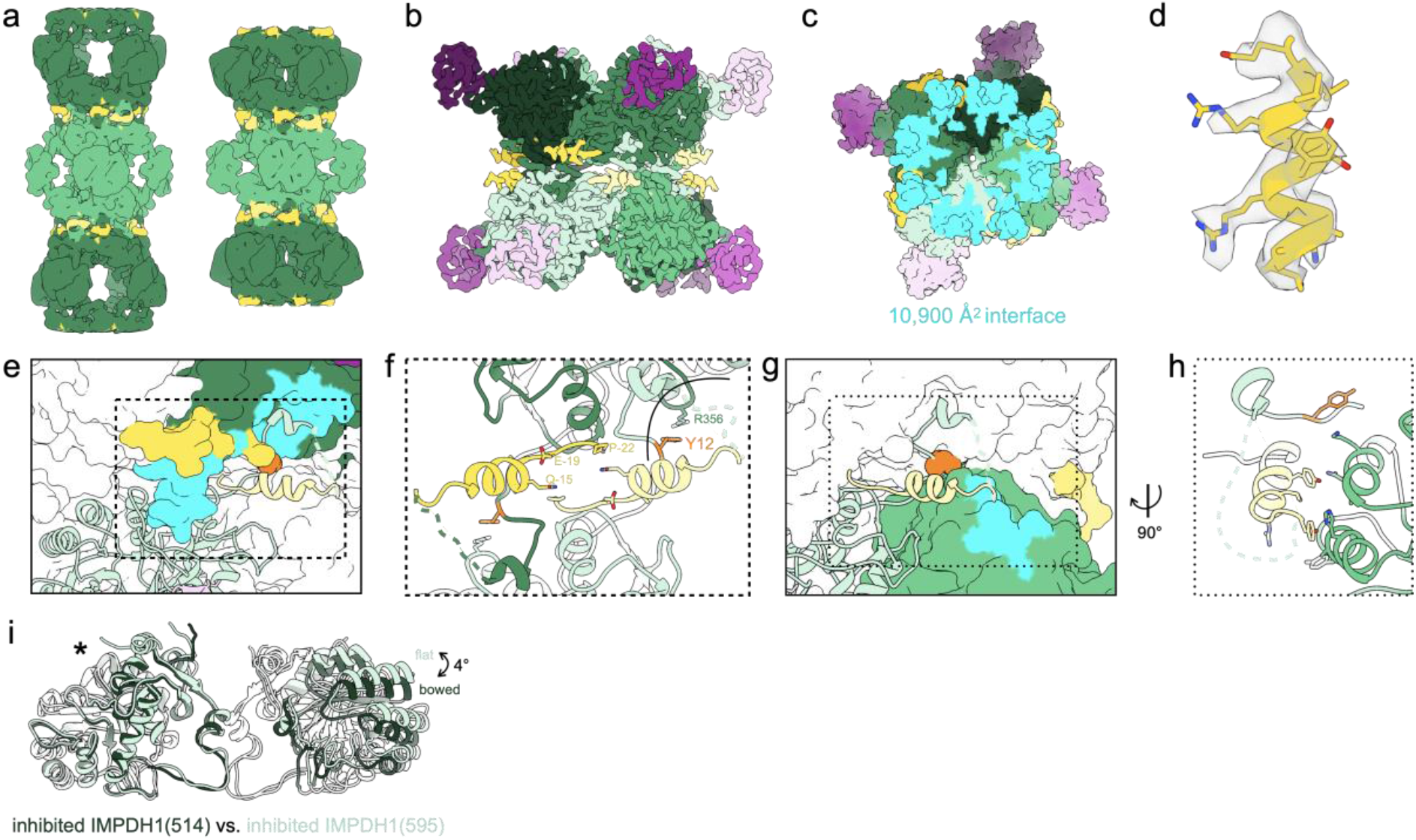
IMPDH1 retinal variant (595) constrains filament architecture. **a**, Low-pass filtered cryo-EM reconstruction of the IMPDH1(595) active filament (bound to ATP) and inhibited filament (bound to GTP, ATP, IMP, NAD+). **b-k**, Inhibited IMPDH1(595) filament (bound to GTP, ATP, IMP, NAD+). **b**, Interface-focused cryo-EM reconstruction. 8 monomers are colored with catalytic domain (green), regulatory domain (pink), and variant helix (yellow). **c**, View of the top of an octamer from inside the filament. The surface area buried by the octamer interface is in aqua with the indicated total buried surface area. (Surface representation of the atomic model at the assembly interface, with buried residues in cyan) **d**, Additional N-term helix residues −18 to −7 in yellow in density. **e**, surface representation of interface with one monomer in ribbon (light green). Additional N-term helix in yellow. The monomer it contacts across the octamer interface is colored dark green/pink with residues the ribbon monomer contacts in aqua. Y12 is shown in orange spheres. **f**, Close-up view of the interface contacts. **g**, Surface representation of interface with one monomer in ribbon (light green). Additional N-term helix in yellow. The neighbor monomer in the tetramer is colored green with residues the ribbon monomer contacts in aqua. Y12 is shown in orange spheres. **h**, Close-up view of the new N-terminal helix contacts with the adjacent monomer in the tetramer. **i**, Comparison of the catalytic tetramers of inhibited IMPDH1(514) GTP/IMP/NAD+ (dark green) to inhibited IMPDH1(595) GTP/ATP/IMP/NAD+ (light green). Aligned on monomers with asterisk.

We next solved structures of IMPDH1(595) in an inhibited filament at 3.7 Å resolution by cryo-EM. Unlike the canonical enzyme, IMPDH1(595) is maintained in the large interface and flat tetramer conformation in the compressed, GTP-bound state. The overall interface and position of the helix from the N-terminal extension are very similar to the active filament described above (1.5 Å RMSD among C*α*s of all eight catalytic domains at the interface when aligned on a single chain). Thus, the role of the N-terminal extension appears to be stabilizing the large interface, and through that stabilizing the flat enzyme conformation that is less sensitive to GTP inhibition. This explains why the effect of the N-terminal extension is dependent on filament assembly (Fig. 5e).

We then wondered how the C-terminal extension contributes to decreased GTP sensitivity. So, we solved a 3.6 Å cryo-EM structure of inhibited IMPDH1(546), which only has the C-terminal extension (Supp. Fig. 6a-6f). In the inhibited conformation, bound to GTP/ATP/IMP/NAD+, IMPDH1(546) assembles with the small interface and in a bowed conformation, very similar to canonical IMPDH1 (Supp. Fig. 6g) In all other structures of inhibited IMPDH, the conformation of the tetramer in the flat/partially inhibited state or bowed/fully inhibited state has explained whether it is more or less sensitive to GTP inhibition. Even though IMPDH1(546) is less sensitive to GTP inhibition than the canonical variant (Fig. 5c), its tetramer is still in the bent/fully inhibited state (Supp. Fig. 6g). Therefore, we suspect the C-terminal extension prevents full octamer compression to resist GTP inhibition. In inhibited canonical IMPDH1, Arg512 and Glu510 near the C-terminus make ionic interactions with residues in protomers on the opposite face of the octamer, stabilizing the compressed conformation. In IMPDH1(546) these residues change to Thr510 and Leu512 (Supp. Fig. 7). Disrupting these interactions likely disfavors octamer compression, potentially explaining why IMPDH1(546) resists GTP inhibition. A second possibility is that the remaining 31 residues that are not resolved in any of our IMPDH1(546) are highly flexible, and their presence near the core of the enzyme may sterically hinder compression.

### Retinopathy mutations fall into two functional classes

Previous studies have looked at the effect of RP mutations only in canonical IMPDH1, where there is no effect on substrate kinetics, but a subset of mutations disrupt GTP regulation^11^. Since disease only occurs in the retina, where there is no expression of canonical IMPDH1^21^, and given the large differences in GTP regulation we observed with splice variants, we wondered whether the mutations had specific effects in the variants. We tested the mutant forms of canonical and retinal IMPDH1 variants for all nine known retinitis pigmentosa (RP) mutations. Our findings for canonical IMPDH1 confirm the previous results^11^. We repeated these experiments with the retinal variants and found the effects of each mutation were similar in the variants and canonical IMPDH1, with several mutations affecting GTP regulation. Thus, in IMPDH1 splice variants, we can describe two classes of disease mutants. Mutations that are insensitive to GTP inhibition we describe as Class I which consists of five mutations around the 3^rd^ allosteric site that is specific for GTP (Fig. 7a–d, Supp. Table 4). Class I mutations are N198K, R231P, R224P, D226N, K238E. Class II mutations are located at four positions more distal to the allosteric sites and had a GTP inhibition response nearly identical to wildtype (Fig. 7e–g, Supp. Table 4). Class II mutations include R105W, T116M, V268I, and H372P. The similar effects of disease mutants in all variants of IMPDH1 suggests that they likely have the same effect at the enzyme level across all tissues expressing IMPDH1.

**Fig 7.**
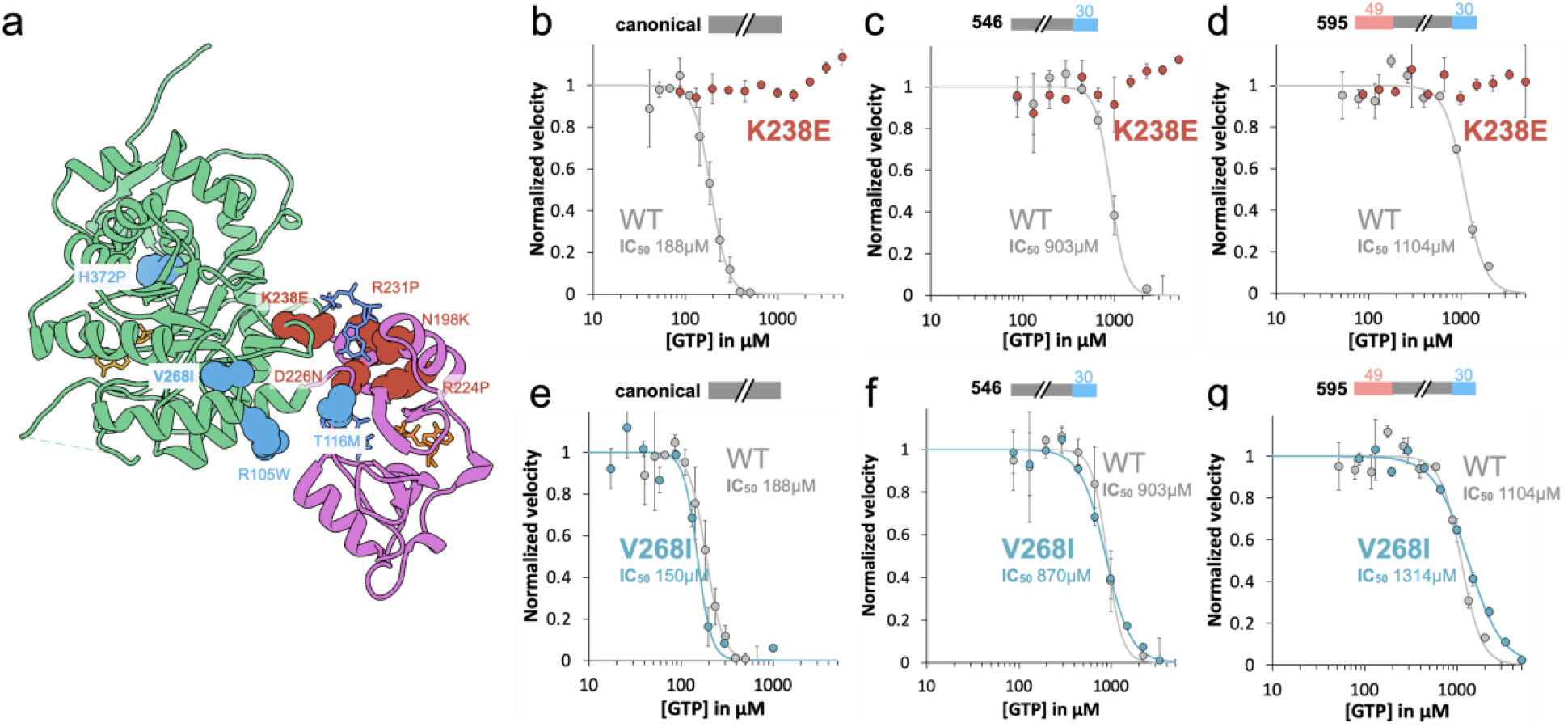
IMPDH1 retinopathy mutations fall into two classes. **a**, IMPDH1 with catalytic domain (green), regulatory domain (pink), NAD+ (gold), GTP (blue), ATP (orange). Class I (defective in GTP regulation) residues colored coral. Class II residues (normal GTP regulation) colored teal. **b-g**, GTP inhibition curves of IMPDH1 retinopathy mutant variants (colored lines) compared to IMPDH1 WT (gray lines). Reactions performed with 1 μM protein, 1 mM ATP, 1 mM IMP, 300 μM NAD+, and varying GTP.

Class II mutations do not have an effect on biochemical activity, so we wondered if the reason they cause disease might be instead due to an effect on filament assembly. We performed negative stain EM in the presence of ATP, substrates, and inhibitory concentrations of GTP (Supp. Fig. 7). Under this condition, all three WT variants assemble filaments of compressed/inhibited octamers. Four of the five Class I mutants form filaments in all variants, but are made up of extended/flexible octamers, which agrees with their inability to be inhibited by GTP. The only Class I mutation that is different is R224P. IMPDH1(514)-R224P forms compressed filaments similar to WT, whereas R224P in both retinal variants does not assemble into filaments (Supp. Fig. 7). In this condition, the Class II mutations form filaments that are indistinguishable from WT in negative stain EM. The lack of obvious in vitro biochemical or structural phenotypes for Class II mutations suggests they may lead to misregulation in vivo that is dependent on other cellular factors.

## DISCUSSION

IMPDH ultrastructure assembly has been observed in many cell types and in vitro assembly of the isoform IMPDH2 has been thoroughly characterized^2,10,33,37^. Despite the essential housekeeping role of IMPDH1 in most cell types and mutations linked to blindness in humans, until now the molecular mechanisms and biochemical consequence of IMPDH1 filament assembly had not yet been described.

Here, we showed that canonical IMPDH1 assembles filaments in both active and inhibited conformations (Figs. 2–4, 8a,b). Other well-characterized filament-forming metabolic enzymes like IMPDH2 and CTPS2 also switch enzyme conformations in the polymer, and constraints imposed by fixed assembly contacts give rise to filament-dependent changes in allosteric regulation^10,38^ (Fig. 8b). IMPDH1, on the other hand, changes both enzyme conformation and the nature of assembly contacts in transitioning between activity states. Thus, filament assembly of canonical IMPDH1 does not impose conformational constraints, explaining why we did not observe any differences in allosteric regulation in the filament compared to free enzyme (Fig. 2d).

**Fig 8.**
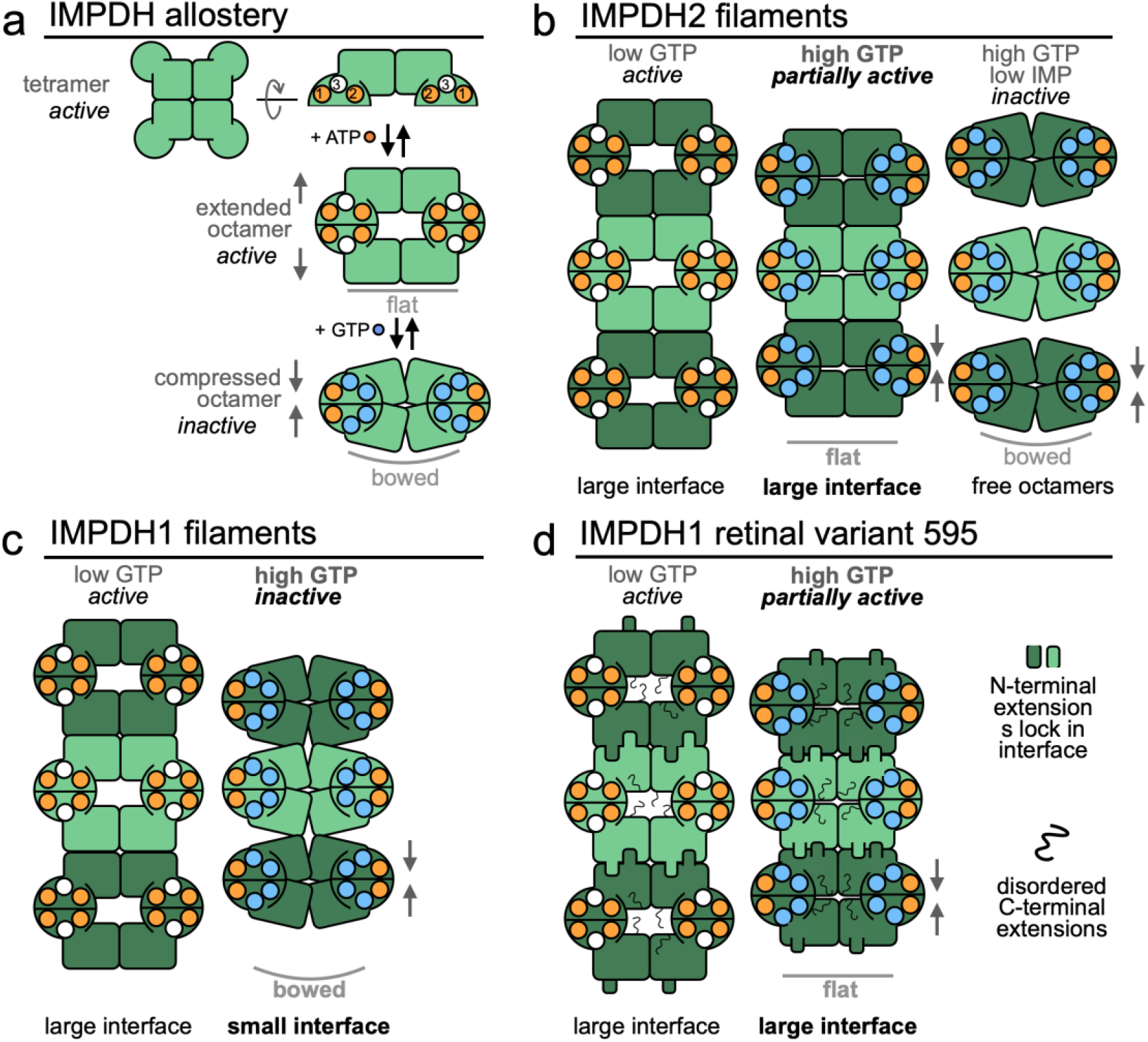
Model of IMPDH1 assembly and filament role in regulation. **a**, ATP binding to sites 1 and 2 promotes formation of an extended/active octamer where the tetramer is in the flat conformation. GTP binding to sites 2 and 3 promotes formation of a compressed/inhibited octamer that prefers the bowed tetramer conformation. **b**, In the presence of ATP, IMPDH2 assembles extended/active filaments. Binding of GTP leads to the assembly of partially inhibited/compressed filaments where the tetramer is in a flat conformation. In the presence of very high GTP, the tetramer enters the complete inhibited/bowed conformation which promotes disassembly of filament into free octamers. **c**, For canonical IMPDH1, binding of ATP drives assembly of a filament composed of extended/active octamers. In the presence of GTP, canonical IMPDH1 assembles into a filament with the small interface made of fully inhibited/compressed octamers that have the bowed tetramer conformation. **d**, For IMPDH1 retinal variant 595, the binding of ATP drives assembly of a filament composed of extended/active octamers. Binding of GTP drives assembly of a filament composed of partially inhibited/compressed octamers that have the strained tetramer confirmation. In both filaments, the N-terminal extension adds buried surface area to the large interface and the C-terminal extension is disordered.

This raises the question of the physiological function of canonical IMPDH1 filaments. Although we show that IMPDH1 specifically assembles filaments in cells in response to inhibitors (Fig. 2b, 8c), how cellular filaments might influence metabolic flux is unclear. One possibility is that IMPDH1 filaments play some role other than direct tuning of enzyme activity, such as signaling or scaffolding of other metabolic enzymes^39,40^. Another possibility is that other cellular factors shift the balance between the active and inhibited filament conformations, for example through post-translational modifications or protein-protein interactions. There is precedent for the latter in the enzyme acetyl-CoA carboxylase (ACC), which assembles active filaments of one architecture, but binding to the regulator breast cancer type 1 susceptibility protein (BRCA1) stabilizes a different filament architecture with inactive enzyme^41^. Similarly, interactions with regulatory proteins may preferentially stabilize one of the IMPDH1 filament states. Indeed, in immunofluorescence experiments using antibodies that recognize both isoforms, IMPDH filaments co-localize with other enzymes in purine biosynthesis^40^, with the regulatory protein Ankyrin Repeat Domain 9 (ANKRD9)^42^, and with morphologically similar CTP synthase ultrastructures^39^. Future studies to identify interactors and to probe the functional consequences of those interactions on IMPDH1 regulation will provide valuable insight into the role of canonical IMPDH1 filament assembly in a physiological context.

This work also sheds light on the role of IMPDH1 filaments in regulating enzyme activity in the retina. The retina has extremely high and specific purine nucleotide demands: maintenance of ion gradients across photoreceptor membranes consumes as many as 10^8^ molecules of ATP per second per cell and cGMP is essential for signaling in phototransduction^24–26^. IMPDH1 is the only isoform expressed in photoreceptors^21^, where it plays a critical role in balancing adenine and guanine nucleotide pools. Two splice variants that add residues to the N- and C-termini of IMPDH1 are predominant in the retina^21^, and we found that both human splice variants are less sensitive to GTP inhibition (Fig. 5c,d).

The N- and C-terminal retinal splice variant extensions independently increase the IC_50_ for GTP, consistent with the need for IMPDH1 to meet high guanine nucleotide demand in the retina. Two structural rearrangements must occur for IMPDH to be fully inhibited: the tetramer is in a bowed conformation and the octamer is fully compressed. We find that each retinal variant extension prevents one of these. The N-terminal extension effect is completely dependent on the ability of the protein to form filaments (Fig. 5e), as it tunes filament assembly so the inhibited protein assembles into the large interface filament that can only accommodate the flat/partially active tetramer. This prevents transition to the small filament interface, so that at high GTP concentrations the filament remains in a partially active compressed and flat conformation (Fig. 8d). The C-terminal extension, on the other hand, appears to destabilize the compressed conformation, either through specific residue changes at the compressed interface or through steric interference of the disordered region (Supp. Fig. 7). The overall effect of the splice variants is to increase IMDPH1 activity at high GTP concentrations, consistent with the high demand for guanine nucleotides in the retina. Meeting this demand with IMDPH1 splice variants instead of IMPDH2 may provide unique regulatory advantages, through protein-protein interactions with, or post translational modification of, the flexible N- and C-terminal extensions.

Some retinopathy mutations in IMPDH1 disrupt GDP feedback inhibition^11^. Why these mutations lead to tissue-specific disease remains unclear, although the fact that the retina only expresses IMPDH1 may make it particularly sensitive to perturbations in the enzyme. We found that disease-associated IMPDH1 mutants fall into two classes: Class I are clustered near GTP allosteric sites and completely disrupt GTP inhibition, while Class II are away from the allosteric sites and have no effect on GTP inhibition. It seems likely that Class I mutations lead to photoreceptor death because they break feedback inhibition and imbalance nucleotide pools, which has been shown to lead to photoreceptor death^43^. Our in vitro characterization of Class II mutations showed that most of them are indistinguishable from wildtype in terms of substrate kinetics and GTP regulation. This finding highlights the need to study these mutations in the complex photoreceptor environment, where IMPDH1 is the only isoform. Understanding the retinopathy mutants in the physiological context of the retina will be necessary to demonstrate the molecular mechanisms of disease.

Many metabolic regulatory enzymes self-assemble into filamentous polymers. In most cases filament assembly serves as an additional layer of allosteric regulation, taking advantage of existing allostery but imposing constraints on accessibility of different conformations ^44,45^. For example, assembly of human CTPS1 into filaments stabilizes a conformation with higher specific activity to increase flux^46^, while CTPS2 assembles filaments that couple structural transitions to increase cooperativity of enzyme regulation ^38^. Yeast glucokinase 1 was recently shown to assemble filaments that inactivate the enzyme, providing a mechanism to reduce overinvestment in early steps of glycolysis on sudden transition to nutrient rich environments ^47^. Acetyl coA carboxylate filaments regulated activity by locking the enzyme into an active state or, in the presence of a regulatory binding partner, into an alternative, inactive assembly ^48^. Here, we have shown that another well-established means of metabolic regulation—tissue-specific splice variants—can add an additional layer of allosteric regulation on top of filament assembly to finely tune complex enzyme regulation (Fig. 8).

## Supporting information

All Supplemental

## ACKNOWLEDGEMENTS

The authors thank the Arnold and Mabel Beckman Cryo-EM Center at the University of Washington for electron microscope use. We thank John Calise for valuable feedback on the manuscript. This work was supported by the US National Institutes of Health (R01GM118396 and R21EY031546 and S10OD032290 to JMK, F31EY030732 and T32GM008268 to ALB).

## Contributions

A.L.B. performed kinetics assays, protein purification, cryo-EM data collection and image processing, and structure analysis. C.N. optimized kinetics experiment design. M.S. developed cryo-EM processing approaches and processed cryo-EM data. D.F.-L. produced WT retinal protein used for cryo-EM. J.Q. optimized cryo-EM sample preparation. R.M.B. and J.R.P. conceptualized experiments. A.L.B. and J.M.K designed experiments, performed data analysis, and interpretation and wrote the manuscript.

## MATERIALS & METHODS

### Recombinant IMPDH expression and purification

Purified IMPDH protein was prepared as described previously^17^. BL21 (DE3) *E. coli* transformed with a pSMT3-Kan vector expressing N-terminal 6xHis-SMT3/SUMO-tagged IMPDH were cultured in Luria broth at 37°C until reaching an OD600 of 0.9 then induced with 1mM IPTG for 4 hr at 30°C and pelleted. The remainder of the purification was performed at 4°C. Pellets were resuspended in lysis buffer (50 mM KPO_4_, 300 mM KCl, 10 mM imidazole, 800 mM urea, pH 8) and lysed with an Emulsiflex-05 homogenizer. Lysate was cleared by centrifugation and SUMO-tagged IMPDH chromatographically purified with HisTrap FF columns (GE Healthcare Life Sciences) and an Äkta Start chromatography system. After on-column washing with lysis buffer and elution (50 mM KPO_4_, 300 mM KCl, 500 mM imidazole, pH 8), peak fractions were treated with 1 mg ULP1 protease^49^ per 100 mg IMPDH for 1 hour at 4°C, followed by the addition of 1 mM dithiothreitol (DTT) and 800 mM urea. Protein was then concentrated using a 30,000 MWCO Amicon filter and subjected to size-exclusion chromatography using Äkta Pure system and a Superose 6 column pre-equilibrated in filtration buffer (20 mM HEPES, 100 mM KCl, 800 mM urea, 1 mM DTT, pH 8). Peak fractions were concentrated using a 10,000 MWCO Amicon filter then flash-frozen in liquid nitrogen and stored at −80°C.

### IMPDH activity assays

Protein aliquots were diluted in activity buffer (20 mM HEPES, 100 mM KCl, 1 mM DTT, pH 7.0) and pre-treated with varying concentrations of ATP, GTP, and IMP for 30 minutes at 20°C in 96 well UV transparent plates (Corning model 3635). Reactions (100 μL total) were initiated by addition of varying concentrations of NAD+. NADH production was measured by optical absorbance (340 nm) in real-time using a Varioskan Lux microplate reader (Thermo Scientific) at 25°C, 1 measurement/min, for 15 minutes; absorbance was correlated with NADH concentration using a standard curve. Specific activity was calculated by linear interpretation of the reaction slope for a 4 minute window beginning 1 minute after reaction initiation. All data points reported are an average of 3 measurements from the same protein preparation. Error bars are standard deviation.

### Negatively stained electron microscopy

Protein preparations were applied to glow-discharged continuous carbon EM grids and negatively stained with 2% uranyl formate. Grids were imaged by transmission electron microscopy using an FEI Morgagni at 100kV acceleration voltage and a Gatan Orius CCD. Micrographs were collected at a nominal 22,000x magnification (pixel size 3.9 Å).

### Electron cryo-microscopy sample preparation and data collection

Protein preparations were applied to glow-discharged C-flat holey carbon EM grids (Protochips), blotted, and plunge-frozen in liquid ethane using a Vitrobot plunging apparatus (FEI) at 4°C, 100% relative humidity. High-throughput data collection was performed using an FEI Titan Krios transmission electron microscope operating at 300 kV (equipped with a Gatan image filter (GIF) and post-GIF Gatan K2 or K3 Summit direct electron detector) and an FEI Glacios (equipped with a Gatan K2 Summit direct electron detector) both using the Leginon software package^50^.

### Electron cryo-microscopy image processing

Movies were collected in super-resolution mode, then aligned and corrected for beam-induced motion using Motioncor2, with 2X Fourier binning and dose compensation applied during motion correction^50,51^. CTF was estimated using GCTF^52^. Relion 3.1 was used for all subsequent image processing ^53,54^. Each dataset was individually processed but using approximately the same previously published pipeline (Johnson and Kollman 2020), with some variations from dataset to dataset.

First, for most datasets, autopicking templates and initial 3D references maps were prepared by manually picking and extracting boxed particles from a small subset of micrographs and classifying/refining in 2D and 3D. For a few data sets, Cryosparc Live^55^ was used for initial particle selection and 2D classification. These particle coordinates were imported into Relion for 3D refinement. For these initial 3D refinements, a featureless, soft-edged cylinder was used as a refinement template of filaments. Because IMPDH filament segments possess D4 point-group symmetry, two different locations along filaments may be used as symmetry origins: the centers of canonical octamer segments, or the centers of the assembly interface between segments. For the filament datasets, we prepared and used auto-picking templates centered on the filament assembly interface. Due to the expected flexibility of filaments, helical segments were processed as single particles, and at no point was helical symmetry applied during image processing. After template-based autopicking of each complete dataset, picked particles were boxed and extracted from micrographs, and subjected to hierarchical 2D classification to select the best-resolved classes. These selected particles were then auto-refined in 3D as a single class with D4 symmetry applied.

To improve resolution, partial signal subtraction was performed at this stage using a mask that left only the central eight catalytic domains of the filament assembly interface, and then subtracting the poorly resolved Bateman domains and neighboring segments, which often improved resolution after subsequent auto-refinement. Per-particle defocus and per-micrograph astigmatism were then optimized using CTF refinement followed by particle polishing, which generally improved resolution further (Supp. Table 5).

### Model building and refinement

Initial templates for model building were prepared from hIMPDH2 extended (PDB 6u8N) and compressed octamers (PDB 6u9o) with amino acid mutations to hIMPDH1 sequence made in Coot ^56^. Where the N-terminus (−22-12) location differed from IMPDH2, it was modeled by hand. After rigid-body fitting of templates into the cryo-EM densities using UCSF Chimera, repeated cycles of manual fitting with Coot and automated fitting with phenix.real_space_refine (employing rigid-body refinement, NCS constraints, gradient-driven minimization and simulated annealing) ^56–58^. Data collection parameters and refinement statistics are summarized in Supplemental Table 5. Figures were prepared with UCSF Chimera ^57^.

The sizes of interacting surfaces between IMPDH protomers were calculated using the PDBePISA server ^59^.

### Cell culture and transfection

HEK293 cells were grown in DMEM/10% fetal bovine serum (FBS)/1% L-glutamine on 6-well dishes with coverslips in each dish to 50% confluency. They were transfected with 40 μl Lipofectamine L-2000 (ThermoFisher Cat #11668030) and 16 μg of pcDNA3.1 plasmid with either IMPDH1-WT or IMPDH1-Y12A for 6 hours and then the media was changed to either new DMEM/10%FBS/1%L-glutamine or DMEM/10%FBS/1%L-glutamine with 10 μM mycophenolic acid (MPA). 24 hours later, the cells were fixed with 4% formaldehyde/PBS for 20 minutes and immunofluorescence was performed on the coverslips.

#### Immunofluorescence

The coverslips with the transfected HEK293 cells were blocked with 2%BSA/PBS and stained with anti-myc antibody 9e10 DSHB (deposited in the Developmental Studies Hybridoma Bank by Bishop, J.M.) diluted 1:50 in 2% bovine serum albumin (BSA)/PBS for 1 hour at room temperature. Then, cells were incubated with Alexa 488 secondary goat anti-mouse (Invitrogen Cat # A-11001) diluted 1:200 in 2%BSA/PBS for 30 minutes at room temperature. Finally, nuclei were stained with DAPI for 10 minutes at room temperature and coverslips were mounted with Vectashield mounting medium (Vector Laboratories). The cells were imaged on a Nikon Eclipse TE2000-U with a 40x Nikon plan fluorescence objective and pictures were taken with Ocular QImaging software Version 1.1 using a QImaging Retiga R1 CCD camera.

### Data availability

The coordinates are deposited in the Protein Data Bank with PDB IDs 7RER (interface-centered active IMPDH1(514)), 7RES (octamer-centered active IMPDH1(514)), 7RFE (interface-centered inhibited IMPDH1(514)), 7RFG (octamer-centered inhibited IMPDH1(514)), 7RGL (interface-centered active IMPDH1(546)), 7RGM (octamer-centered active IMPDH1(546)), 7RGI (interface-centered inhibited IMPDH1(546)), 7RGQ (octamer-centered inhibited IMPDH1(546)), 7RFF (interface-centered active IMPDH1(595)), 7RFH (octamer-centered active IMPDH1(595)), 7RFI (interface-centered inhibited IMPDH1(595)), 7RGD (octamer-centered inhibited IMPDH1(595). The cryo-EM maps are deposited in the Electron Microscopy Data Bank with IDs EMD-24437 (interface-centered active IMPDH1(514)), EMD-24438 (octamer-centered active IMPDH1(514)), EMD-24439 (interface-centered inhibited IMPDH1(514)), EMD-24441 (octamer-centered inhibited IMPDH1(514)), EMD-24451 (interface-centered active IMPDH1(546)), EMD-24452 (octamer-centered active IMPDH1(546)), EMD-24450 (interface-centered inhibited IMPDH1(546)), EMD-24454 (octamer-centered inhibited IMPDH1(546)), EMD-24440 (interface-centered active IMPDH1(595)), EMD-24442 (octamer-centered active IMPDH1(595)), EMD-24443 (interface-centered inhibited IMPDH1(595)), EMD-24448 (octamer-centered inhibited IMPDH1(595).

