## Supplementary material for "IMPDH1 retinal variants control filament architecture to tune allosteric regulation": All Supplemental

### Supplementary Figures

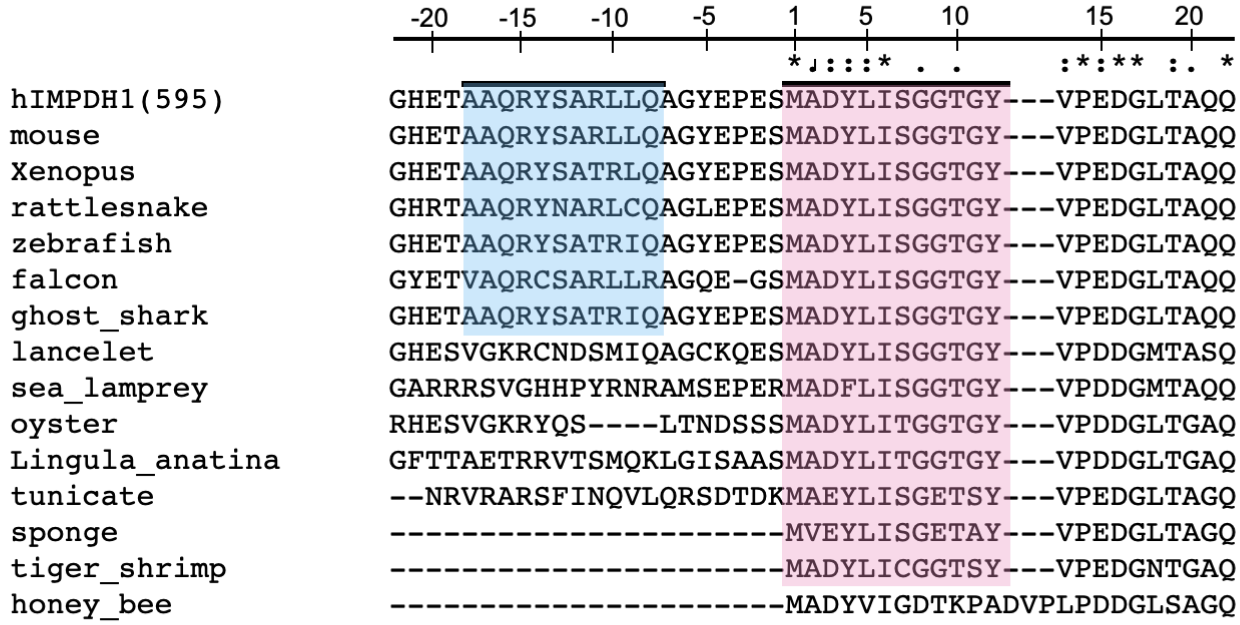

**Supplemental Fig 1. Sequence Alignment**
Evolutionary conservation of the helix in the N-terminus of the longer retinal splice variant (blue) and the first 12 canonical residues particularly tyrosine 12 (pink).

| | | IMPDH1 | IMPDH2 |
|---|---|---|---|
| IMP | **Vmax** (µM NADH/min/µM IMPDH) | 3.5 | 2.8 |
| | **Km** (µM) | 60 | 38 |
| | **Hill** | 3.5 | 3.0 |
| NAD | | **IMPDH1** | **IMPDH2** |
| | **Vmax** (µM NADH/min/µM IMPDH) | 3.4 | 5.3 |
| | **Km** (µM) | 16.3 | 27 |
| | **Hill** | 2.6 | 1.4 |

**Supplemental Table 1. IMPDH1 and IMPDH2 have similar kinetics**
(A-E) Table comparing $V_{max}$, $K_m$, and Hill coefficients for IMP and NAD+ for IMPDH1 and IMPDH2. Reactions performed with 1 µM protein, 1 mM ATP, 1 mM or varying IMP, and 300 µM or varying NAD+.

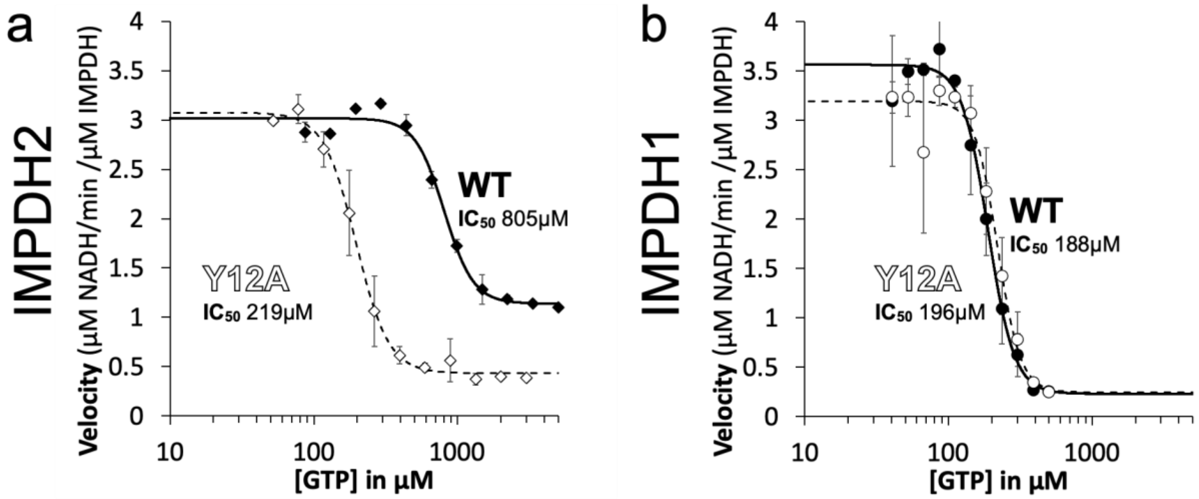

**Supplemental Fig 2. Inhibited IMPDH2-WT filament resists GTP inhibition**

**a-b**, GTP inhibition curves of IMPDH2 or IMPDH1 WT (solid line) and the respective non-assembly Y12A protein (dashed line). Reactions performed with 1 µM protein, 1 mM ATP, 1 mM IMP, 300 µM NAD+, and varying GTP.

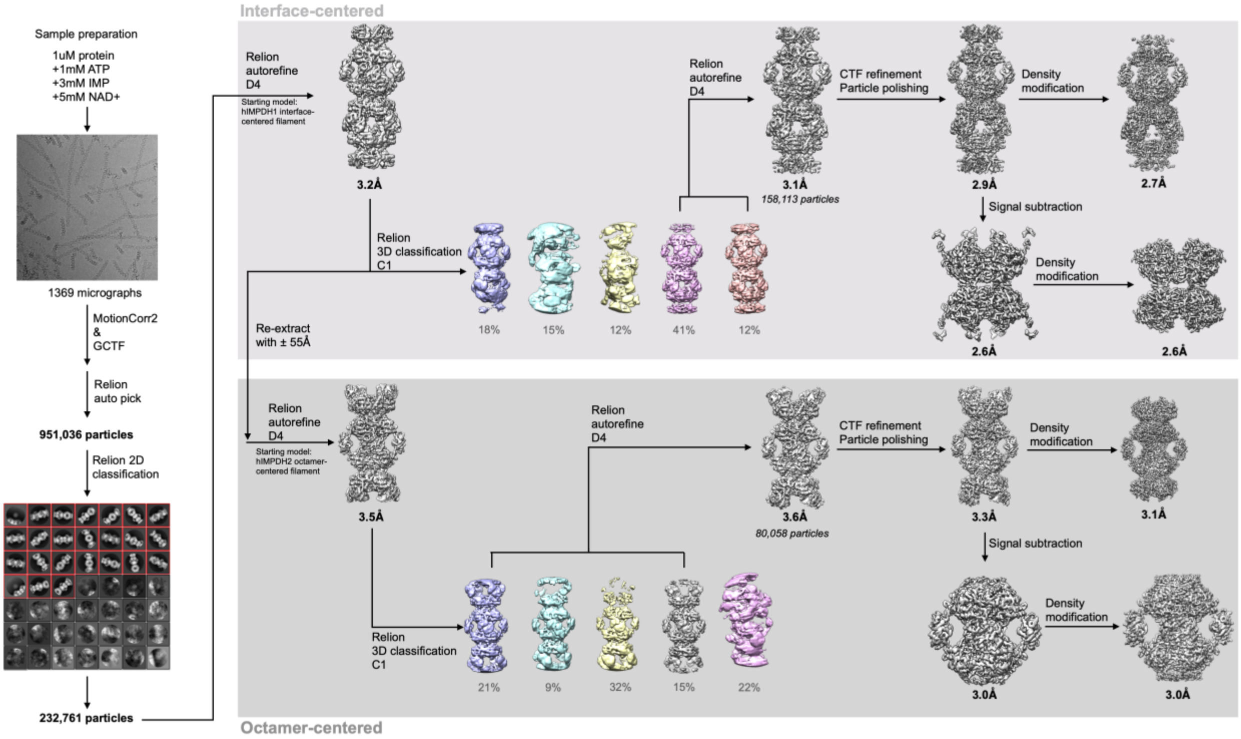

**Supplemental Fig. 3**
Flow chart summarizing data processing strategy for IMPDH1 + ATP/IMP/NAD+

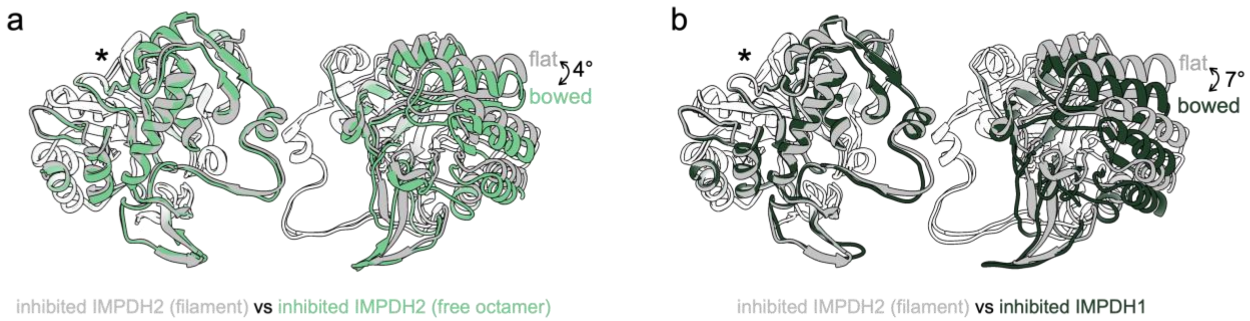

**Supplemental Fig 4. Inhibited IMPDH1-WT tetramer is in a bowed conformation**

**a**, Comparison of the catalytic tetramers of inhibited IMPDH2 filament (gray; 6u8s) to inhibited IMPDH2 free octamer (6uaj). Aligned on monomers with asterisk, other monomer pair has an alpha carbon RMSD of 2.1 Å. **b**, Comparison of the catalytic tetramers of inhibited IMPDH2 filament (gray; 6u8s) to inhibited IMPDH1 filament. Aligned on monomers with asterisk, other monomer pair has an alpha carbon RMSD of 3.7 Å.

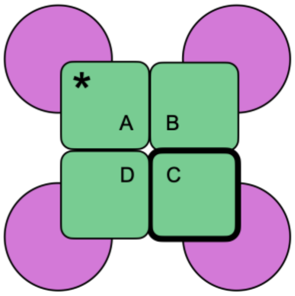

| | | IMPDH2 | | IMPDH1(514) | | IMPDH1(546) | | IMPDH1(595) | |
|---|---|---|---|---|---|---|---|---|---|
| | | Active large interface | Inhibited large interface | Active large interface | Inhibited small interface | Active large interface | Inhibited small interface | Active large interface | Inhibited large interface |
| IMPDH2 | Active large interface | — | 0.061 | 0.878 | 3.314 | 2.145 | 3.246 | 2.712 | 0.693 |
| | Inhibited large interface | — | — | 1.266 | 1.721 | 2.371 | 2.371 | 2.595 | 0.816 |
| IMPDH1(514) | Active large interface | — | — | — | 3.787 | 1.705 | 3.7 | 2.249 | 1.065 |
| | Inhibited small interface | — | — | — | — | 5.046 | 1.368 | 5.506 | 2.208 |
| IMPDH1(546) | Active large interface | — | — | — | — | — | 4.723 | 0.554 | 1.698 |
| | Inhibited small interface | — | — | — | — | — | — | 5.174 | 1.924 |
| IMPDH1(595) | Active large interface | — | — | — | — | — | — | — | 0.560 |
| | Inhibited large interface | — | — | — | — | — | — | — | — |

**Supplemental Table 2. RMSD between catalytic domains of models**
Top-down view of tetramer with catalytic domains (green) and regulatory domains (pink). Models are aligned on catalytic domain with asterisk and alpha carbon RMSD is calculated by catalytic domain outlined in bold. IMPDH2 active is 6u8n, IMPDH2 inactive is 6u8s RMSD to determine how tetramer flexing (bent/flat) compares.

(video submitted as separate file)

**Supplemental Video 1. Comparison between large and small interface IMPDH1 filaments**
Morph comparison between large interface of ATP/IMP/NAD+ IMPDH1 filament to small interface in GTP/ATP/IMP IMPDH1 filament.

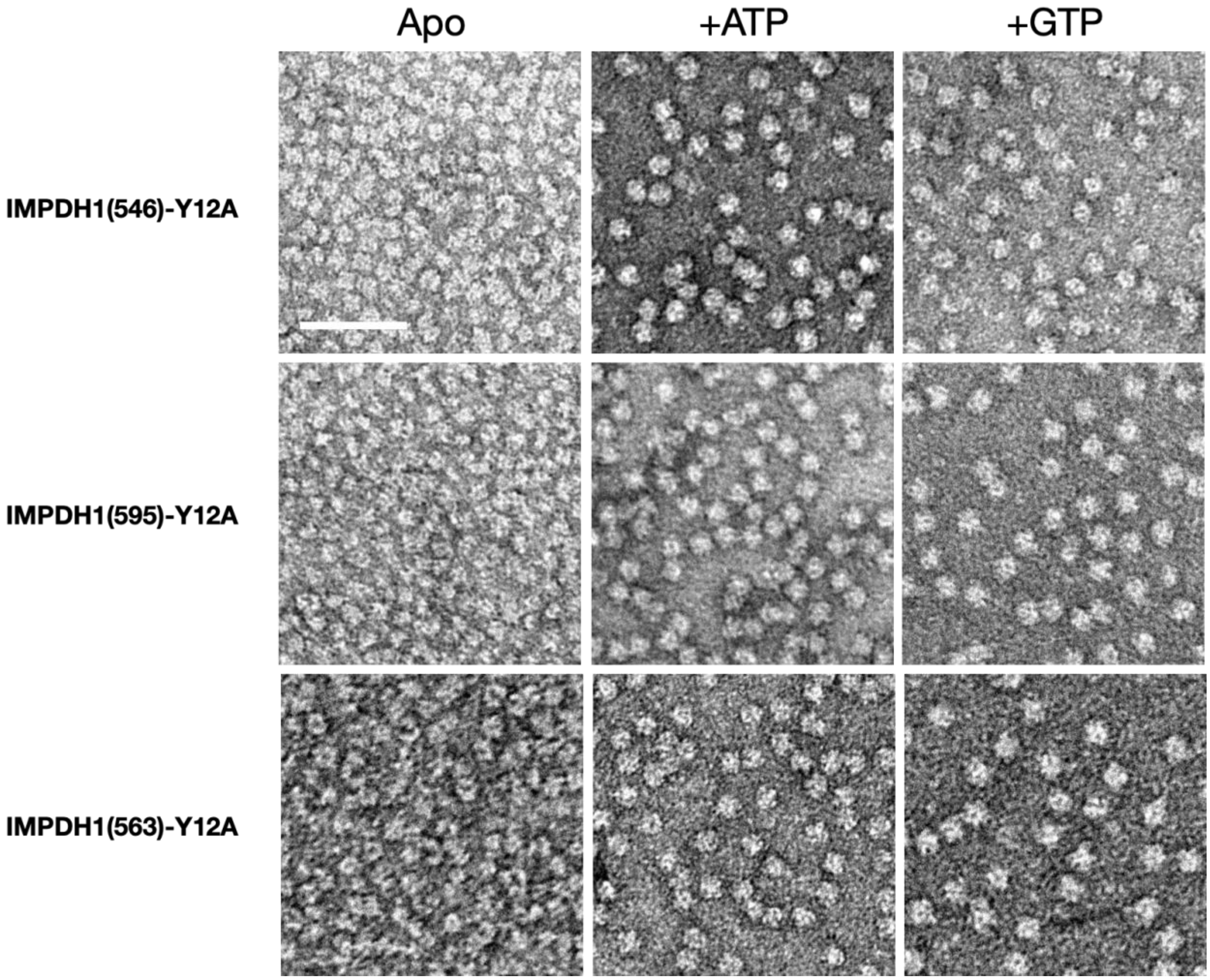

**Supplemental Fig 5. Y12A non-assembly mutations prevents assembly in IMPDH1 variants**
Negative stain EM of purified human IMPDH1. Non-assembly mutation Y12A breaks both ATP- and GTP-dependent assembly. Scale bar 100 nm. Reactions performed with 1 µM protein, 1 mM ATP if used, 1 mM GTP if used.

| | WT | Y12A |
|---|---|---|
| **IMPDH2** | 805 | 219 |
| **IMPDH1(514)** | 188 | 196 |
| **IMPDH1(546)** | 903 | 585 |
| **IMPDH1(563)** | 632 | 151 |
| **IMPDH1(595)** | 1104 | 550 |

**Supplemental Table 3. IMPDH1 and IMPDH2 $IC_{50}$ for GTP in μM**

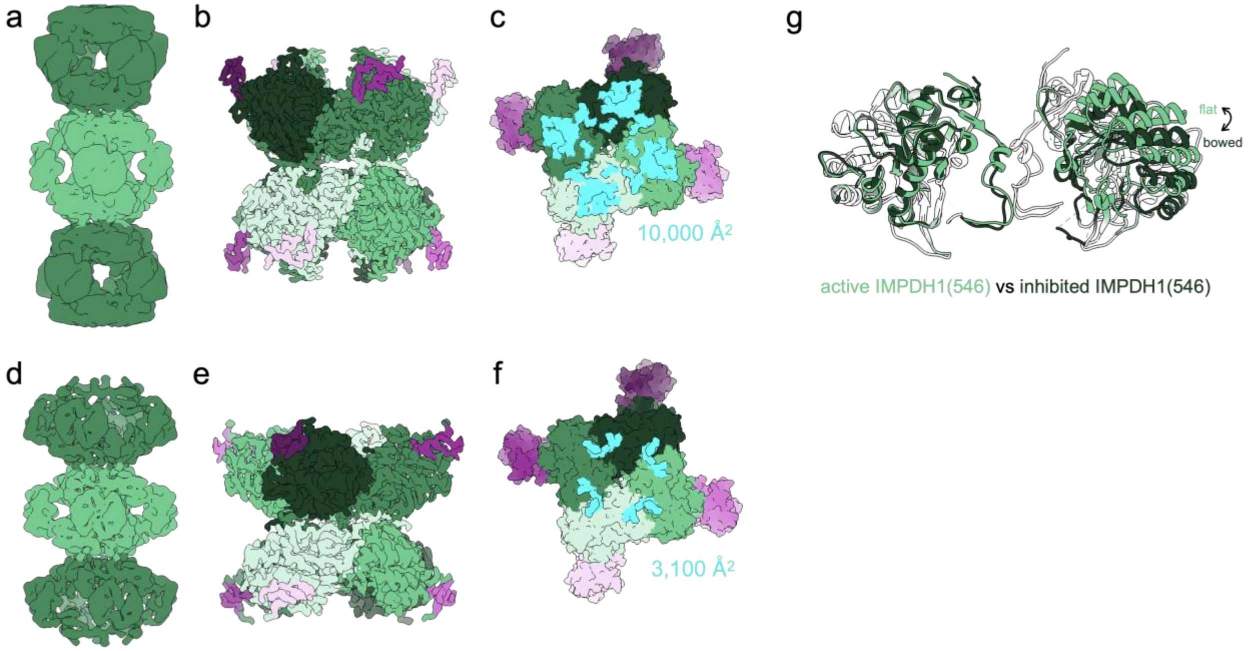

**Supplemental Fig 6. IMPDH1 retinal variant (546) is similar to canonical IMPDH1**

**a-c**, Active IMPDH1(546) filament bound to ATP, IMP, NAD+. **a**, Low-pass filtered cryo-EM reconstruction **b**, Interface-focused cryo-EM reconstruction. 8 monomers are colored by catalytic domain (green) and regulatory domain (pink). **c**, View of the top of an octamer from inside the filament. The surface area buried by the octamer interface is in aqua with the indicated total buried surface area. (Surface representation of the atomic model at the assembly interface, with buried residues in cyan). **d-f**, Inhibited IMPDH1(546) filament bound to GTP, ATP, IMP, NAD+. **d**, Low-pass filtered cryo-EM reconstruction **e**, Interface-focused cryo-EM reconstruction. 8 monomers are colored by catalytic domain (green) and regulatory domain (pink). **f**, View of the top of an octamer from inside the filament. The surface area buried by the octamer interface is in aqua with the indicated total buried surface area. (Surface representation of the atomic model at the assembly interface, with buried residues in cyan).

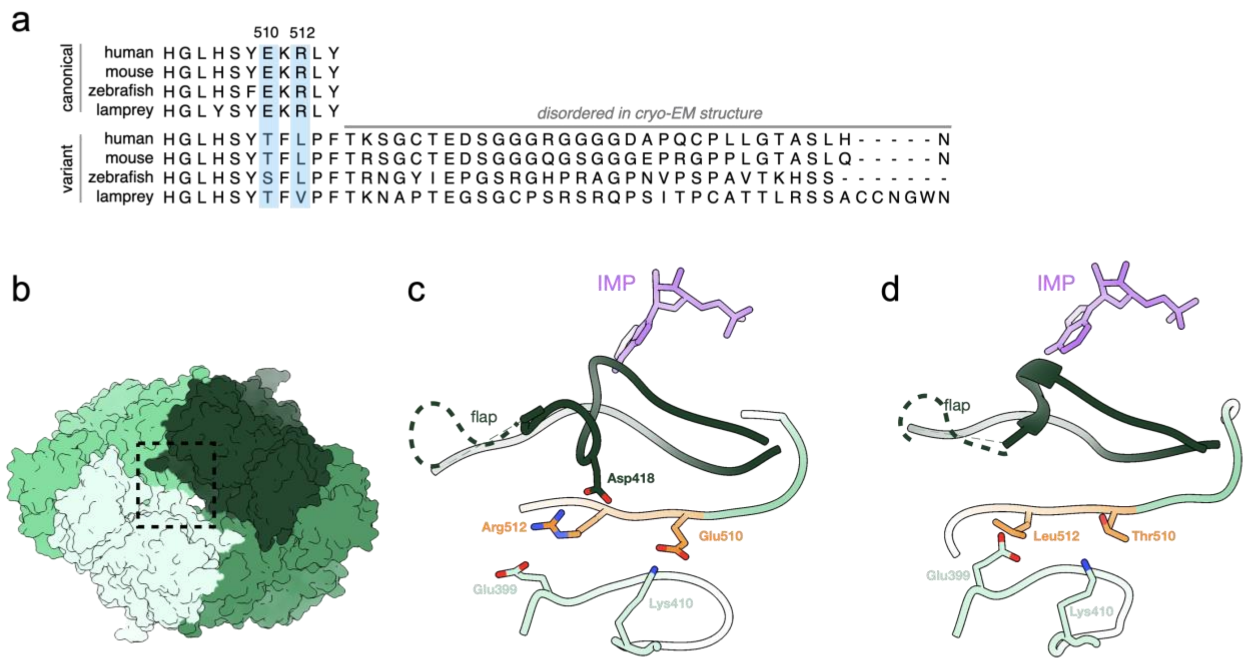

**Supplemental Fig 7. IMPDH1 Retinal Variant C-term disrupts interactions**

**a**, evolutionary conservation of the C-terminus in canonical IMPDH1 and both retinal splice variants. **b**, Surface representation of octamer side view. Dotted box indicates the region shown in **c-d**. **c-d**, Each chain is a different color green, C-term residues 510-512 in orange, and IMP in purple. **c,** inhibited canonical IMPDH1. **d**, Inhibited retinal variant IMPDH1(546).

**a**

| | | WT | R105W | T116M | N198K | R224P | D226N | R231P | K238E | V268I | H372P |
|---|---|---|---|---|---|---|---|---|---|---|---|
| 514 | Vmax | 3.4 | 3.7 | 3.4 | 3.4 | 3.2 | 3.4 | 4.2 | 5.3 | 4.6 | 4.4 |
| | Km | 16.3 | 15.5 | 9.3 | 10.8 | 9.9 | 8.8 | 12.9 | 20.7 | 16.9 | 16.2 |
| | Hill | 2.6 | 3.1 | 1.7 | 3.2 | 1.9 | 1.7 | 2.1 | 2.7 | 2.4 | 3.2 |
| 546 | | WT | R105W | T116M | N198K | R224P | D226N | R231P | K238E | V268I | H372P |
| | Vmax | 5.5 | 5.3 | 4.2 | 5.1 | 6.4 | 5.1 | 5.8 | 5.4 | 6.0 | 8.8 |
| | Km | 32.9 | 22.8 | 19.5 | 31.0 | 31.5 | 26.2 | 31.9 | 22.5 | 24.9 | 46.9 |
| | Hill | 1.9 | 1.1 | 1.3 | 1.0 | 1.0 | 1.1 | 0.9 | 1.0 | 0.9 | 1.1 |
| 595 | | WT | R105W | T116M | N198K | R224P | D226N | R231P | K238E | V268I | H372P |
| | Vmax | 5.2 | 5.3 | 4.0 | 4.8 | 1.5 | 5.5 | 6.8 | 26.9 | 5.1 | 7.4 |
| | Km | 30.4 | 32.6 | 28.8 | 36.7 | 28.1 | 23.1 | 39.6 | 1.6 | 36.7 | 66.2 |
| | Hill | 1.2 | 1.0 | 1.1 | 0.6 | 0.7 | 0.9 | 1.1 | 0.9 | 1.3 | 1.3 |

**b**

| | | WT | R105W | T116M | N198K | R224P | D226N | R231P | K238E | V268I | H372P |
|---|---|---|---|---|---|---|---|---|---|---|---|
| 514 | Vmax | 3.5 | 4.2 | 2.5 | 3.0 | 3.1 | 2.6 | 3.5 | 5.0 | 3.3 | 3.5 |
| | Km | 59.5 | 13.6 | 28.0 | 19.9 | 12.3 | 10.0 | 20.2 | 17.9 | 25.4 | 25.0 |
| | Hill | 3.5 | 2.7 | 2.7 | 2.1 | 2.0 | 1.9 | 2.2 | 2.0 | 2.5 | 3.0 |
| 546 | | WT | R105W | T116M | N198K | R224P | D226N | R231P | K238E | V268I | H372P |
| | Vmax | 4.8 | 5.3 | 4.7 | 4.7 | 4.5 | 4.6 | 4.5 | 4.5 | 4.8 | 6.1 |
| | Km | 33.0 | 23.3 | 34.4 | 34.4 | 27.6 | 22.6 | 17.9 | 26.8 | 39.2 | 45.1 |
| | Hill | 1.3 | 1.0 | 0.8 | 0.8 | 1.3 | 1.1 | 1.2 | 1.5 | 1.4 | 1.3 |
| 595 | | WT | R105W | T116M | N198K | R224P | D226N | R231P | K238E | V268I | H372P |
| | Vmax | 5.2 | 4.9 | 3.6 | 3.7 | 5.0 | 4.6 | 6.0 | 6.2 | 5.0 | 7.1 |
| | Km | 27.9 | 20.4 | 23.4 | 18.9 | 22.3 | 22.6 | 26.0 | 35.8 | 20.3 | 28.6 |
| | Hill | 1.4 | 1.5 | 1.6 | 1.2 | 1.1 | 1.1 | 1.3 | 1.5 | 1.3 | 1.5 |

**c**

| | | WT | R105W | T116M | N198K | R224P | D226N | R231P | K238E | V268I | H372P |
|---|---|---|---|---|---|---|---|---|---|---|---|
| 514 | Vmax | 3.6 | 4.1 | 2.7 | 3.0 | 2.8 | 2.2 | 3.2 | 4.5 | 3.1 | 3.3 |
| | IC50 | 187.8 | 15.9 | 132.4 | N/A | N/A | 134.0 | N/A | N/A | 150.8 | 151.2 |
| | Hill | 4.7 | 25.2 | 3.0 | N/A | N/A | 32.1 | N/A | N/A | 5.7 | 2.6 |
| 546 | | WT | R105W | T116M | N198K | R224P | D226N | R231P | K238E | V268I | H372P |
| | Vmax | 4.4 | 4.8 | 4.4 | 3.9 | 5.2 | 4.5 | 3.9 | 4.5 | 5.3 | 7.3 |
| | IC50 | 903.4 | 759.6 | 888.3 | N/A | N/A | N/A | N/A | N/A | 870.9 | 630.8 |
| | Hill | 5.3 | 2.7 | 3.8 | N/A | N/A | N/A | N/A | N/A | 3.2 | 5.4 |
| 595 | | WT | R105W | T116M | N198K | R224P | D226N | R231P | K238E | V268I | H372P |
| | Vmax | 4.5 | 4.9 | 4.1 | 3.6 | 4.9 | 4.5 | 5.0 | 6.0 | 5.2 | 7.4 |
| | IC50 | 1104.0 | 1274.4 | 1760.5 | N/A | N/A | N/A | N/A | N/A | 1313.9 | 1843.0 |
| | Hill | 3.9 | 2.4 | 3.5 | N/A | N/A | N/A | N/A | N/A | 2.4 | 1.8 |

**Supp Table 4. IMPDH1 RP mutations do not change NAD+ or IMP kinetics in any variants**
**A-c**, $V_{max}$ (µM NADH/min/µM IMPDH), $K_m$ (µM), and Hill coefficient for NAD+ for WT and RP mutants in all variants. **a**, For NAD+. Reactions performed with 1 µM protein, 1 mM ATP, 1 mM IMP, and varying NAD+. **b**, For IMP. Reactions performed with 1 µM protein, 1 mM ATP, 300 µM NAD+, and varying IMP. **c**, For GTP. Reactions performed with 1 µM protein, 1 mM ATP, 1 mM IMP, 300 µM NAD+, and varying GTP.

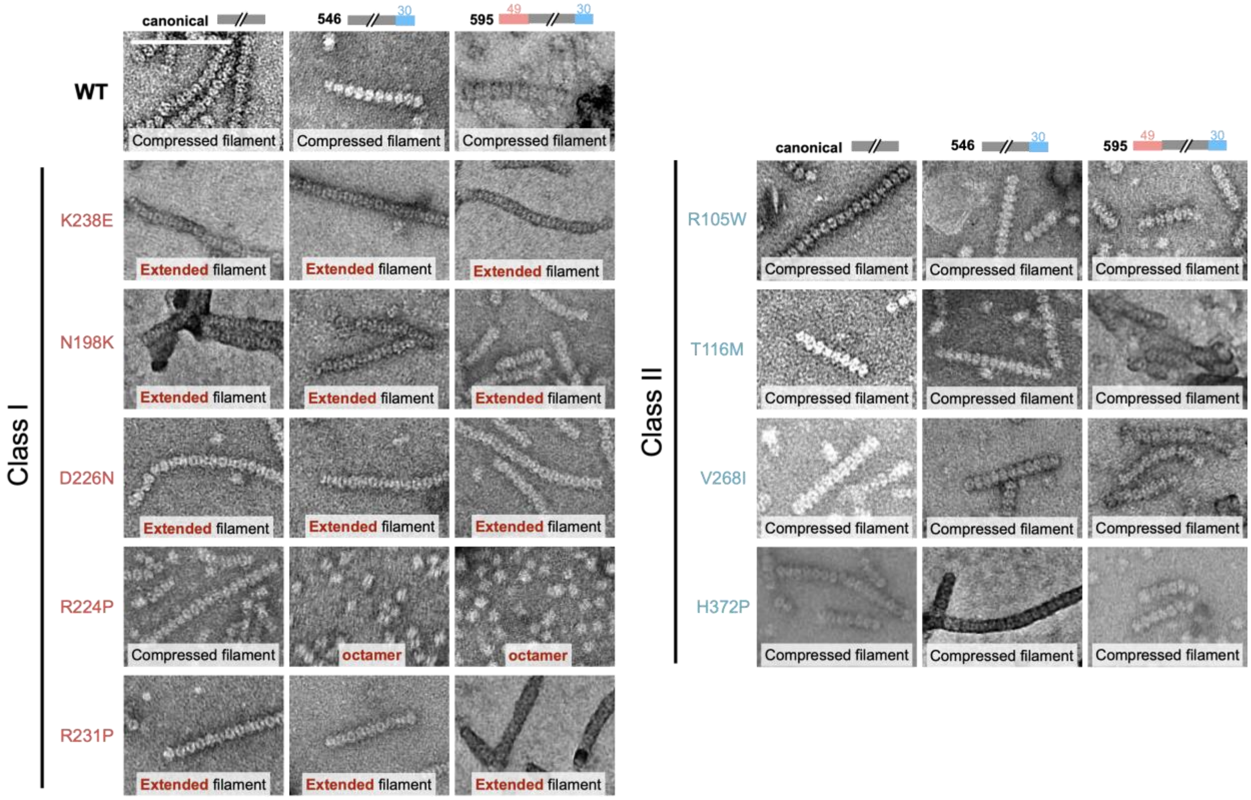

**Supplemental Fig 7. IMPDH1 disease mutants have a variety of assembly phenotypes**
Negative stain EM of purified human IMPDH1. Scale bar 100 nm. Reactions performed with 1 µM protein, 1 mM ATP, 5 mM GTP, 3mM IMP, 5mM NAD+.

| | IMPDH1(514) | | | | IMPDH1(546) | | | | IMPDH1(595) | | | |
|---|---|---|---|---|---|---|---|---|---|---|---|---|
| | Active (ATP, IMP, NAD+) | | Inhibited (GTP, ATP, IMP) | | Active (ATP, IMP, NAD+) | | Inhibited (GTP, ATP, IMP, NAD+) | | Active (ATP) | | Inhibited (GTP, ATP, IMP, NAD+) | |
| **Data collection** | | | | | | | | | | | | |
| Voltage (kV) | 300kV | | 300kV | | 300kV | | 200kV | | 300kV | | 300kV | |
| Defocus range (-µm) | 1.5-3.5 | | 0.5-1.8 | | 0.6-1.8 | | 0.5-3.2 | | 0.6-2.3 | | 0.5-2.5 | |
| Pixel size (Å) | 0.525 | | 0.525 | | 0.4215 | | 1.16 | | 0.525 | | 0.4215 | |
| **Reconstruction** | | | | | | | | | | | | |
| | Interface | Octamer | Interface | Octamer | Interface | Octamer | Interface | Octamer | Interface | Octamer | Interface | Octamer |
| Pixel size (Å) | 1.05 | 1.05 | 1.05 | 1.05 | 0.843 | 0.843 | 1.16 | 1.16 | 1.05 | 1.05 | 0.843 | 0.843 |
| Symmetry imposed | D4 | D4 | D4 | D4 | D4 | D4 | D4 | D4 | D4 | D4 | D4 | D4 |
| Particles | 103,809 | 57,965 | 103,500 | 58,000 | 25,366 | 43,406 | 70,031 | 97,369 | 60,919 | 11,476 | 71,792 | 102,543 |
| Map resolution (0.143 FSC, Å) | 2.6 | 3.1 | 2.6 | 3.1 | 2.4 | 2.9 | 4.0 | 3.9 | 2.7 | 3.7 | 2.8 | 3.0 |
| **Refinement** | | | | | | | | | | | | |
| Map sharpening B factor ($Å^2$) | -87 | -102 | -55 | -52 | -38 | -63 | -137 | -133 | -113 | -93 | -57 | -62 |
| Model compostion | | | | | | | | | | | | |
| Ligands | NAD+, IMP | ATP, IMP, NAD+ | GTP, ATP, IMP | GTP, ATP, IMP | IMP, NAD+ | ATP, IMP, NAD+ | IMP, NAD+ | GTP, ATP, IMP, NAD+ | | ATP | GTP, IMP, NAD+ | GTP, ATP, IMP, NAD+ |
| R.m.s deviations | | | | | | | | | | | | |
| Bond lengths (Å) | 0.0111 | 0.0101 | 0.0171 | 0.0064 | 0.0091 | 0.0169 | 0.0091 | 0.0058 | 0.0135 | 0.0118 | 0.0160 | 0.0189 |
| Bond angles (degree) | 0.82 | 0.77 | 1.22 | 0.73 | 0.85 | 1.11 | 0.85 | 0.84 | 0.78 | 1.01 | 1.00 | 1.15 |
| **Validation** | | | | | | | | | | | | |
| MolProbity score | 1.72 | 2.54 | 2.70 | 2.62 | 2.22 | 2.68 | 2.22 | 2.24 | 2.84 | 3.01 | 2.67 | 3.02 |
| Clashscore | 7.77 | 11.52 | 14.10 | 12.66 | 14.84 | 9.58 | 14.84 | 13.28 | 20.85 | 52.61 | 17.75 | 26.95 |
| Poor rotamers (%) | 0 | 5 | 6 | 6 | 0 | 7 | 0 | 0 | 5 | 1 | 4 | 6 |
| **Ramachandran plot** | | | | | | | | | | | | |
| Favored (%) | 96 | 93 | 93 | 94 | 90 | 91 | 90 | 88 | 92 | 75 | 93 | 92 |
| Allowed (%) | 4 | 7 | 6 | 6 | 10 | 9 | 10 | 12 | 8 | 24 | 7 | 8 |
| Outliers (%) | 0 | 0 | 1 | 0 | 0 | 0 | 0 | 0 | 0 | 1 | 0 | 0 |
| **Depostion** | | | | | | | | | | | | |
| EMDB | 24437 | 24438 | 24439 | 24441 | 24451 | 24452 | 24450 | 24454 | 24440 | 24442 | 24443 | 24448 |
| RCSB | 7RER | 7RES | 7RFE | 7RFG | 7RGL | 7RGM | 7RGI | 7RGQ | 7RFF | 7RFH | 7RFI | 7RGD |

**Supplemental Table 5. Cryo-EM Data Collection and Processing Information**
